# Antimicrobial peptides modulate long-term memory

**DOI:** 10.1101/328286

**Authors:** Raquel Barajas Azpeleta, Jianping Wu, Jason Gill, Ryan Welte, Chris Seidel, Kausik Si

**Affiliations:** Stowers Institute for Medical Research, 1000E 50^th^ Street, Kansas City, MO 64110; Department of Molecular and Integrative Physiology, University of Kansas Medical Center, 3901 Rainbow Boulevard, Kansas City, Kansas 66160

**Keywords:** long-term memory, antimicrobial peptides

## Abstract

Antimicrobial peptides act as a host defense mechanism and regulate the commensal microbiome. To obtain a comprehensive view of genes contributing to long-term memory we performed mRNA sequencing from single *Drosophila* heads following behavioral training that produces long-lasting memory. Surprisingly, we find that two immune peptides with antimicrobial activity, Diptericin B and Gram-Negative Bacteria Binding Protein like 3, regulate long-term but not short-term memory or instinctive behavior in *Drosophila*. The cellular requirement of these two peptides is distinct: head fat body for DptB, and neurons for GNBP-like3. That antimicrobial peptides influence memory provides a novel example of the emerging link between the immune and nervous systems and reveals that some immune peptides may have been repurposed in the nervous system.

## Author summary

It is becoming evident that the nervous system and immune system share not only some of the same molecular logic but also the same components. Here, we report a novel and unanticipated example of how immune genes influence nervous system function. During exploring how *Drosophila* form long-lasting memories of certain experiences, we have found that antimicrobial peptides that fight bacteria in the body, are expressed in the head, and control whether an animal would form long-term memory of a food source or a mating partner. Antimicrobial peptides are detected in the brain of many species and has often been associated with dysfunction of the nervous system. This and other recent works, provide an explanation to why antimicrobial peptides may be expressed in the brain: they regulate normal functions of the brain. Both eating, and mating engage the immune system in preparation of exposure to external agents including bacteria. We speculate antimicrobial peptides were upregulated in the body to deal with immune challenges and over evolutionary time some of them are co-adopted to convey specific information about food or mating to the brain.

## Introduction

In most animals modifying behavior based on past experiences is important for survival and reproductive success. To achieve these experience-dependent behavioral modifications, organisms must form memories of specific situations and maintain them to guide future behavior. Given that animals encounter different types of experiences, the resulting memories also vary in nature and duration. Moreover, not only the types of event, but also the internal state of the organism, influences whether an animal will form memory of a given experience, or, if memory is formed, how long it will persist. At molecular level it remains unclear how an animal forms various types of memories with different durations in different context.

The immune system and nervous system rely on their ability to detect and discriminate many cues from the external environment and produce appropriate responses. Similarly, once a cue is encountered, both systems possess the ability to modify their response to the same cue in subsequent encounters. Given the similarity in functional logic, therefore, it is perhaps not surprising that several immune genes also function in the nervous system. One of the earliest example of this is the major histocompatibility complex 1, which is expressed both in the developing and mature nervous system of mice. The MHC1 genes are important for synaptic pruning as well as synaptic plasticity [1]. Likewise, the complement system has been shown to be important for synapse formation, and immune receptors, such as Toll receptors, peptidoglycan pattern recognition receptor (PGRP), or interleukin receptors, are important for synaptic plasticity [2,3,4]. In *Drosophila* immune peptides have been implicated in sleep regulation [5] and nonassociative learning [6].

In course of exploring how animals form long-lasting memories, we discovered, surprisingly, that peptides that are known to be induced in the body upon bacterial infection, are induced in the adult fly head following behavioral training that produces long-term memory. Two of these immune peptides, Diptericin B (DptB) and Gram-Negative Bacteria Binding Protein like 3 (GNBP-like3), are required for efficient long-term memory formation but not for immediate memory. We also find that these peptides attenuate bacterial growth consistent with their posited antimicrobial function. Antimicrobial peptides modulating specific aspects of memory provides a novel example of the emerging link between the immune and nervous systems and leads us to propose that some immune peptides might have been repurposed in the nervous system to “moonlight” as neuromodulators over the course of evolution. It is unclear at this stage how these immune peptides modulate long-term memory.

## Results

### A mRNA sequencing-based screen to identify long-term memory related genes

To identify genes involved in long-term memory, it is common to compare changes in gene expression between trained and untrained animals at a specific time after training. However, training exposes animals to multitude of stimuli, all of which change gene expression, and only a subset of gene expression changes is related to long-term memory *per se*. In addition, animals are continuously responding to a dynamic environment, resulting in differences in gene expression between individuals over time.

To circumvent these problems, and identify genes that specifically regulate long-term memory, we performed mRNA sequencing from individual 4-5 day-old fly heads after training in two distinct memory paradigms (**Fig 1**): male courtship suppression paradigm (MCS), in which a virgin male fly learns to suppress its instinctive courtship behavior after repeated rejection from an unreceptive female fly; and associative appetitive conditioning (AAC), where starved flies learn to associate a specific odor (conditioning stimuli; CS) with a food reward (unconditioning stimuli; US). We used these two paradigms because while both paradigms produce long-term memory, they differ in number of ways: 1) MCS is the modification of an instinctive behavior driven by reproductive urge that requires several hours of training, whereas AAC is a learned behavior driven by hunger that requires 5 minutes of training; 2) while MCS is a single fly behavior, AAC is a group behavior; and 3) while MCS assesses male flies, AAC evaluates male and female flies allowing us to eliminate sex-specific differences. We reasoned a comparative analysis, and identification of the common genes may help isolate genes that are specifically involved in long-term memory, from genes that are involved in other aspects of animal behavior or physiology.

**Figure 1.**
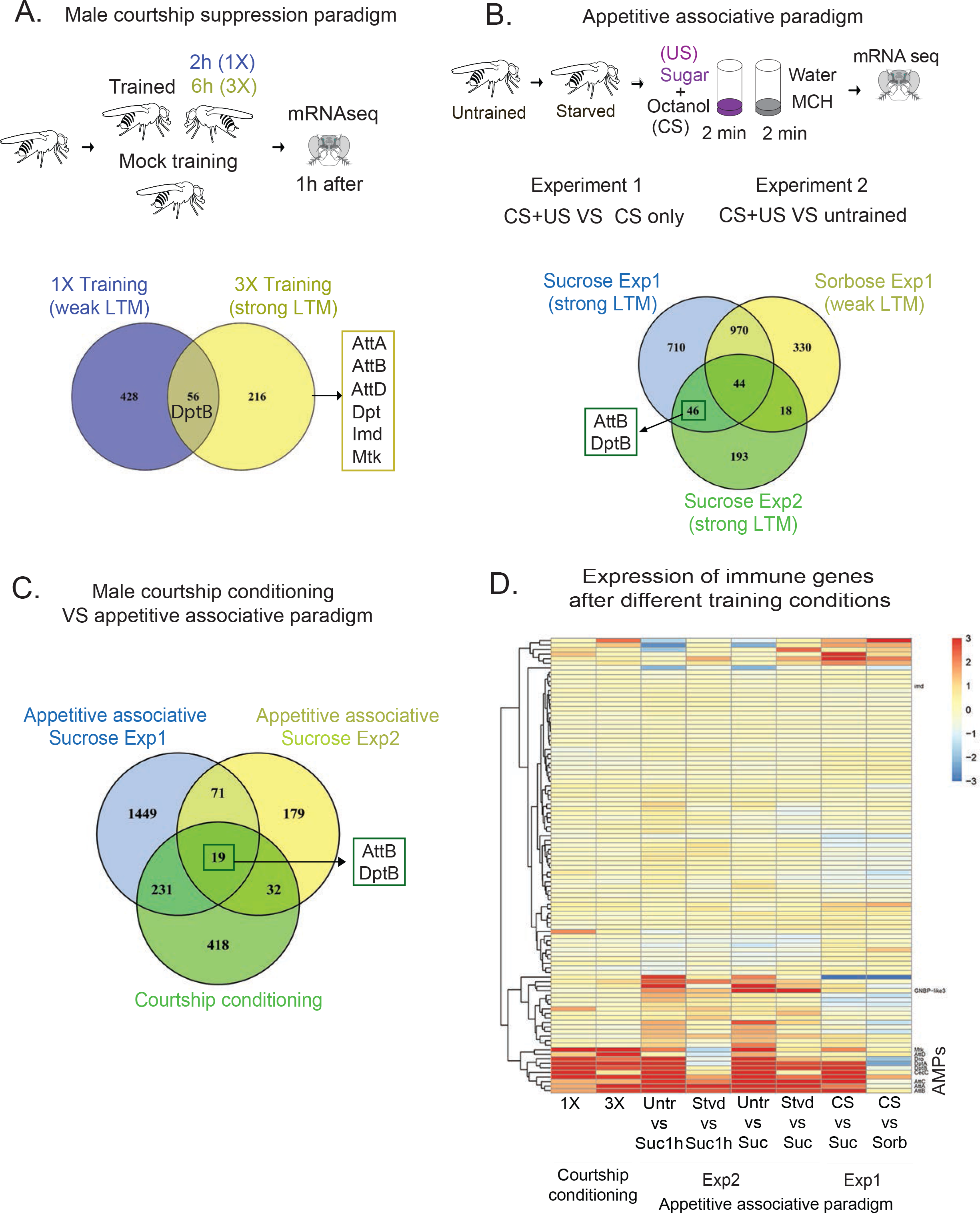
Transcripts of some antimicrobial peptides are upregulated in adult fly head following behavioral training that produces long-term memory. (**A**) Top, schematic representation of male courtship suppression training and mRNA sequencing. Bottom, Venn diagram depicting gene expression changes after 1X (blue), and 3X (yellow) training compared to the mock trained group. The antimicrobial peptide genes that are up-regulated after 3X training are indicated in the box. (**B**) Top, schematic representation of appetitive associative memory paradigm and mRNA sequencing. Bottom, Venn diagram comparing gene expression changes 4 hour after training with sucrose (two independent experiments carried out a month apart) or L-sorbose, compared to the CS only control group (Experiment 1), or the untrained control (Experiment 2). Antimicrobial peptide genes that are changed in both sucrose groups but not in L-sorbose are indicated in the box. (**C**) Venn diagram comparing gene expression changes after male courtship suppression conditioning, and appetitive associative training. (**D**) Heat map showing the expression of immune genes 1 hour or 4 hours after different training conditions. The antimicrobial family is specifically altered in adult head under various behavioral conditions. Att: Attacin, CecC: CecropinC, Dpt: DiptericinA, DptB: DiptericinB, Dro: Drosocin, GNBP-like3: Gram-Negative Binding Protein like3, Imd: Immune deficiency, Mtk: Metchnikowin.

In male courtship suppression a virgin male was exposed to an unreceptive mated female for 2 hours (1X training), or 6 hours (3X training with a gap of 30 minutes between each training session). Single training leads to weak long-term memory, while repeated training results in robust long-term memory [7,8]. We sequenced mRNA from individual virgin male fly heads 1 hour after 1X, or 3X training, or mock trained group as a control (**Fig 1A**). Genes that are changed in the trained group compared to controls are tabulated (**S1 Table**). Seven hundred genes were significantly up or down regulated in the trained groups (padj<0.05) compared to the mock trained control. From those 700 genes, 56 were common to both 1X and 3X training.

In appetitive associative conditioning, starved flies are trained to associate octanol (CS) to sucrose (US). Four hours after training mRNA was sequenced from 4 to 6 individual female fly heads and compared that to age matched untrained fed flies. To distinguish gene expression changes linked to CS-US association from gene expression changes due to starvation, or exposure to odor or sweet sugar, we also performed mRNA sequencing from three control groups; i) starved flies exposed to just octanol (CS only), ii) starved flies trained with octanol and L-sorbose (a sweet, non-nutritious sugar that produces robust short-but weak long-term memory [9,10] (**S1 Fig**) as US, and iii) an independent group of flies trained with sucrose as US after a month, to rule out the variation in gene expression in different population of flies (**S2 Table**). Expression of mRNA that changed only in both sucrose-trained groups compared to naive, CS alone or sorbose trained group, were deemed to be associated with long-term appetitive memory (**Fig 1B**, **S2 Fig** **and S2 Table**). Around 1800 genes were up or down (p<0.05) regulated in the CS+US (sucrose) group, compared to the CS only group. However, when compared to sorbose as US, the number was reduced to ~750 genes. In the second fly population trained one month later, around 300 genes were up or down regulated after training the flies with sucrose compared to the untrained control group. Interestingly, only 46 genes were common in both sucrose experiments, underscoring the variation in gene expression in different population of flies.

### Behavioral training induces DptB expression in the adult *Drosophila* head

We looked for the common genes that were up-or down regulated in both paradigms as candidate longterm memory genes. Expectedly, transcripts of some genes already implicated in memory such as Gclm (Box), Tig (Avgust) [11] were changed following behavioral training (**S3 Fig**). However, to our surprise, a group of immune peptides that are known to be induced in the body upon microbial infection [12,13], such as AttacinB and DptB, were uniquely upregulated in the adult head when male flies are trained to suppress their instinctive courtship behavior, or when starved flies learned to associate odor with sucrose (**Fig 1C**). To verify this unusual observation, we compared our sequencing results with two published datasets that analyzed the change in gene expression at a different time point after 3X MCS training by microarray and mRNA sequencing [14] (**S4 Fig** **and S3-S4 Table**). Although these experiments were carried out in different conditions in different labs, expression of DptB was significantly upregulated in trained animals in both studies. We wondered whether the stress associated with training resulted in a general change in immune gene expression. To this end, we looked at the expression level of all known immune genes in our data sets and observed that a subclass, the immune peptides of antimicrobial family, are consistently altered in adult head under various behavioral conditions (**Fig 1D**).

Nonetheless this unusual observation prompted us to further investigate the specificity of antimicrobial peptide induction under various training conditions. To accomplish this, we used the DptB gene since it consistently changed in all analyses. RT-qPCR (**Fig 2A**) showed that 3X training results in a 10-12-fold increase in DptB mRNA level, compared to mock trained group. Furthermore, induction of DptB was significantly attenuated when the male fly was exposed to a decapitated mated female, instead of a live mated female (**Fig 2A**). This suggests that the experience of active rejection is important for the optimal induction of DptB, and mere exposure to a mated female is not enough. Similarly, using RT-qPCR, we further verified that in appetitive conditioning, training with octanol as CS and sucrose as US also results in DptB mRNA upregulation within one hour after training (**Fig 2B**). Taken together these results suggest that the expression of some bacteria-induced immune peptides, such as DptB, are induced in the adult head when animals are trained to modify their behavior over a long period of time.

**Figure 2.**
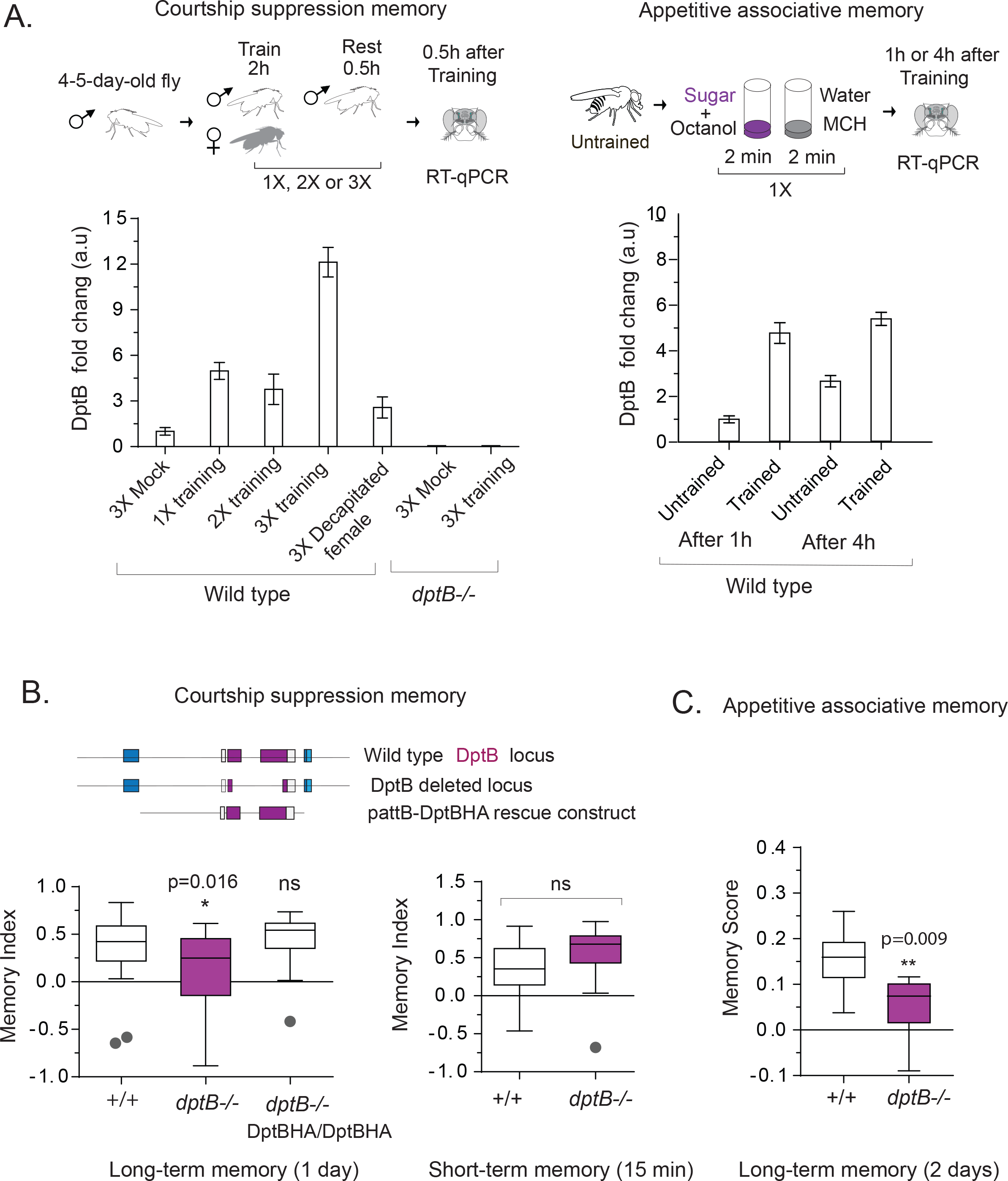
DptB is required for long-term memory. (**A**) DptB expression measured by RT-qPCR after male courtship suppression paradigm: 1X training, 2X training, 3X training, and 3X training with decapitated females (left). DptB expression measured by RT-qPCR 1 hour or 4 hours after associative appetitive memory paradigm (right). The data are mean of three technical repeats. (**B**) Knocking out DptB significantly reduces long-term memory. Long-term memory (24 hours after training) and short-term memory (15 minutes after training) of DptB null flies. Long-term memory deficit of DptB null flies is rescued by a DptB genomic construct. The dots indicate outliers. (**C**) Removal of DptB impairs long-term appetitive memory. The data are plotted as mean ± SEM. Statistical analysis was performed using unpaired two-tailed t-test and (*) P≤ 0.05, (**) P≤0.01, (***) P≤0.001 and (ns) not significant.

### DptB is required for long-term but not for short-term memory or innate behavior

What is the functional relevance, if any, of behavior-dependent increase in antimicrobial peptide (AMP) expression? To this end using Crispr-Cas9 based gene-editing system we deleted the DptB gene. dptB null flies (*dptB*−/−) were viable with no discernable developmental problem. When wild type or *dptB*−/− male flies were trained in courtship suppression paradigm and memory was measured one day after training, *dptB*−/− flies showed a significant reduction (p=0.016) in memory compare to wild type control (**Fig 2B**). This behavioral deficit is rescued by 2 copies of a genomic fragment encompassing the DptB gene. The *dptB*−/− flies also had a significantly (p=0.009) reduced capacity to form long-term appetitive associative memory (**Fig 2C**).

Is DptB affecting memory, or the effect in memory is due to a general disruption in nervous system function? Knocking out DptB gene had no effect in short-term memory (**Fig 2B**), or in innate courtship behavior, such as courtship latency, copulation latency, or duration of copulation (**S5 Fig**). Likewise, the detection of sucrose was unchanged between wild type or mutant flies in a broad concentration (1M-1mM) range (**S5 Fig**). We also tested the effect of removal of DptB in the heat box-paradigm, an operant conditioning paradigm [15] that measures short-term avoidance behavior, locomotion, and sensory perception to noxious stimuli. However, the removal of DptB had no effect in any of these behaviors (**S6 Fig**). Taken together these results suggest DptB plays an important role specifically in animals’ ability to form and/or retain long-term memory.

### DptB is required in the head fat body for long-term memory

Where is DptB made to aid long-term memory? The AMPs are synthesized primarily in the fat bodies, a major secretory tissue that controls metabolism and immune response. In the adults, the fat bodies are found in the abdomen and in the head, surrounding the brain. The head fat body has been implicated in sex-specific and courtship behavior [16,17]. Therefore, we first used an inducible head-fat-body-GAL4 line to express RNAi against the DptB in the adult stage, and measured courtship suppression memory 24 hours after training. We not only analyzed DptB, but also other AMPs whose mRNA level changed following behavioral training as well as some other AMPs whose mRNA expression is not significantly altered in trained flies. As additional control for non-specific effect of activating RNAi pathway in memory, we also expressed dsRNA against luciferase and mCherry. Only the expression of DptB RNAi, no other AMPs or control Luciferase or mCherry RNAi, in head fat body resulted in a significant (p<0.01) reduction of long-term memory (**Fig 3A**). The effect was specific to long-term memory since DptB RNAi had no effect on short-term memory (**Fig 3B**), innate courtship behavior (**S5 Fig**) or short-term operant conditioning or locomotion (**S6 Fig**). Moreover, head fat body appears to be the only relevant source of DptB for behavioral modification, since expression of DptB RNAi in neurons, body fat body, or glial cells, other possible sources of AMPs, had no effect in long-term memory (**Fig 3C**). Taken together these results suggest DptB peptide made in the adult head body influence long-lasting memory.

**Figure 3.**
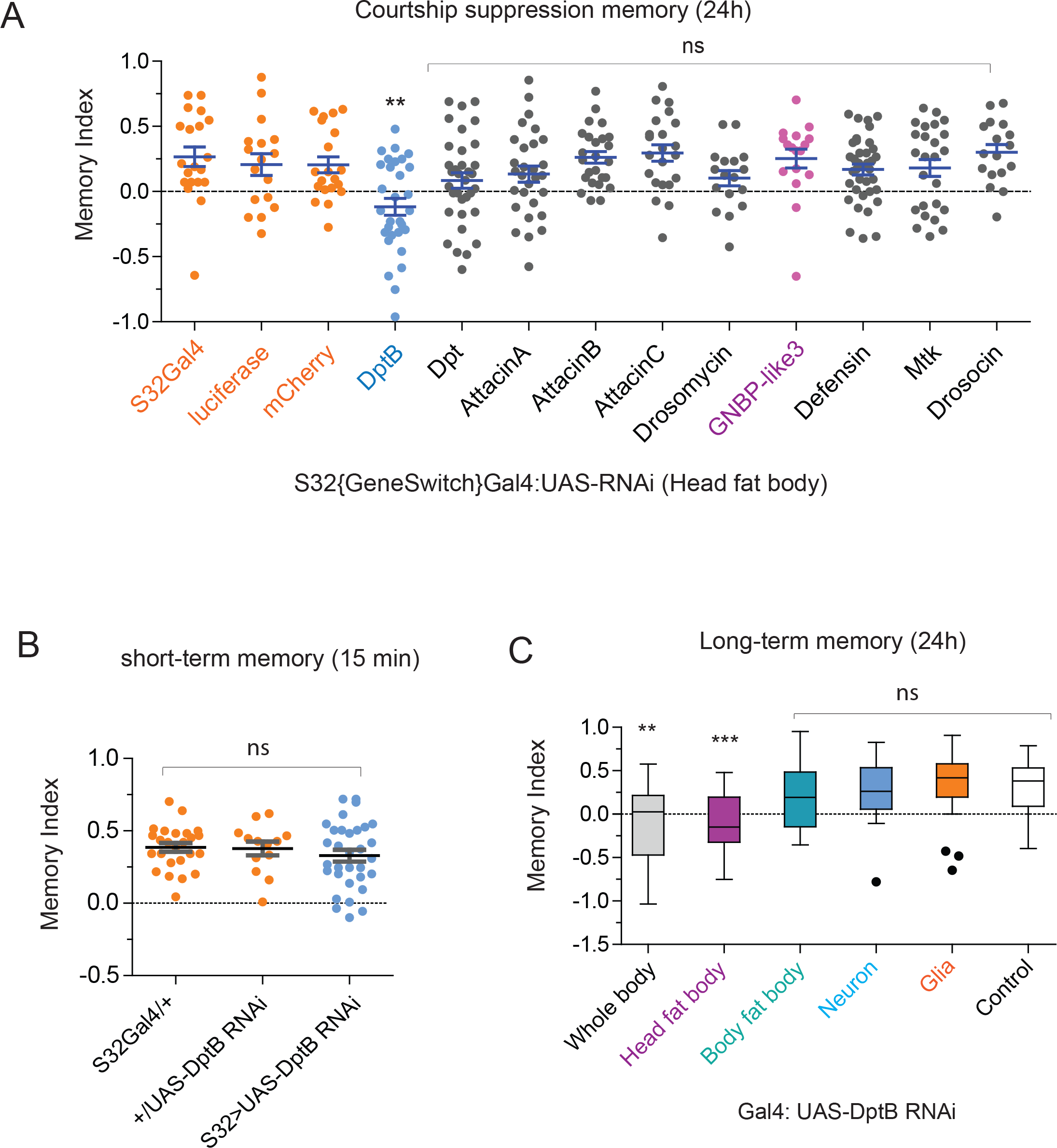
DptB is required in the head fat body for long-term memory. (**A**) Memory index of different antimicrobial peptide RNAis expressed in head fat body 24 hours after male courtship suppression paradigm. Only DptB RNAi flies show a significant reduction in male courtship suppression memory compared to the control groups. (**B**) Short-term male courtship suppression memory is unaffected by DptB knock down. (**C**) Expression of DptB only in head fat body is important for courtship suppression memory. Statistical analysis was performed using one-way ANOVA comparison and (**) P≤0.01, (***) P≤0.001 and (ns) not significant. DptB: DiptericinB, GNBP-like3: Gram-Negative Binding Protein like3.

### GNBP-like 3 is required in neurons for long-term memory

Surprisingly, expression of RNAi in head fat body against some of the other AMPs, such as AttacinB, had no effect on behavior. This suggests that except DptB, other AMPs, although induced, are not required for memory. Alternatively, the tissue source of the other AMPs is different. Based on the observation that in other species immune genes can be expressed in the neurons [18,19], we used a pan-neuronal-GAL4 line, to express RNAi against the AMPs in the nervous system and measured long-term courtship suppression memory (**Fig 4A**). As indicated before the expression of DptB RNAi in neurons had no effect, however, surprisingly, the expression of GNBP-like3 RNAi in neurons significantly (p<0.001) impaired long-term courtship suppression memory (**Fig 4A**). GNBP-like3 is one of the immune peptides whose mRNA level did not significantly change upon behavioral training. The memory phenotype is not likely a non-specific effect of activation of RNAi pathway in neuron, since expression of dsRNA against other genes had no effect in long-term memory (**Fig 4A**) and reduction of GNBP-like3 had no effect in short-term memory (**Fig 4B**) or instinctive behavior of the flies (**S7 Fig**). Likewise, except neurons, expression of GNBP-like3 RNAi just in head fat body, body fat body or glia had no effect in long-term memory (**Fig 4C**). Nonetheless, to ensure that the phenotype is indeed due to loss of GNBP-like3, we generated GNBP-like3 null flies (*gnbp-like3* −/−) using Crispr-Cas9 gene editing system. The *gnbp-like3*−/−flies also had a long-memory deficit (**Fig 4D**).

**Figure 4.**
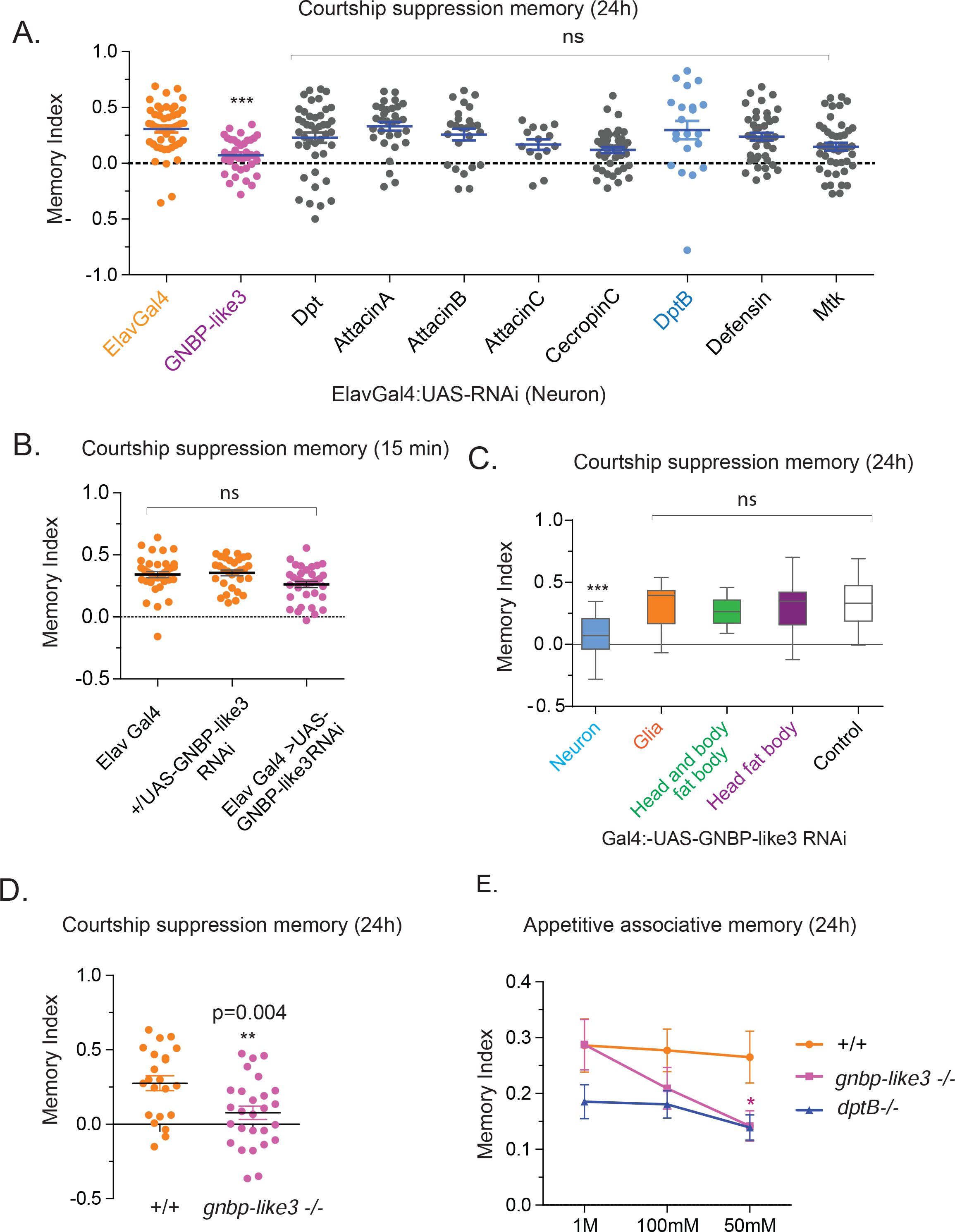
GNBP-like 3 is required in neurons for long-term memory. (**A**) Memory index of different antimicrobial peptide RNAis expressed in neurons 24 hours after male courtship suppression paradigm. The expression of GNBP-like3 RNAi in neurons significantly impaired long-term memory. (**B**) Shortterm male courtship suppression memory is unaffected by GNBP-like3 knock down in neurons. (**C**) Expression of GNBP-like3 only in neurons is important for courtship suppression memory. (**D**) Deletion of GNBP-like3 gene impairs the ability to form long-term associative appetitive memory. (E) 24 h memory of wild type and AMP deficient flies at different sucrose concentration as US. The data are plotted as mean ± SEM. Statistical analysis was performed using one-way ANOVA comparison and (**) P≤0.01, (***) P≤0.001 and (ns) not significant in A. Statistical analysis was performed using unpaired two-tailed t-test and (*) P≤ 0.05, (**) P≤0.01, (***) P≤0.001 and (ns) not significant in B, C, D, and E.

Interestingly, in appetitive-associative memory, *gnbp-like3*−/− did not show any memory loss in higher sucrose concentration (1M sucrose, memory index: Wt, 0.28±0.04, *gnbp-like3*−/−, 0.28± 0.5). However, when the concentration of sucrose was dropped to 50mM, *gnbp-like3*−/−flies had significantly reduced memory (50mM sucrose, memory index: Wt, 0.26±0.46, *gnbp-like3*−/−, 0.14±0.027, p=0.031, student t-test) compared to wild type flies (**Fig 4E**). These observations suggest that the requirement of DptB and GNBP-like3 in appetitive memory may be tuned to the stimulus intensity. Since the sugar concentration in natural food sources are likely to vary [20,21], they may be required for efficient memory formation under varying conditions.

Taken together, these results suggest that DptB and GNBP-like3 serve very specific functions in longterm memory. Surprisingly, even though upregulation of AttacinB was strongly co-related with long-term memory, expression of AttacinB RNAi in either the head fat body or in neurons did not interfere with memory. Either its function is not memory related, or it is required in a specific cell population that we have not been able to interrogate, or the AttacinB RNAi did not perturb its function adequately. Likewise, we can’t rule out the possibility that the lack of phenotypes for other immune peptides may also have resulted from inadequate knockdown by the respective RNAi.

### DptB is expressed in the head fat body, and GNBP-like3 in the central nervous system

We were surprised that tissue requirement of these peptides to influence memory is distinct. We wondered whether this is a simple reflection of their respective expression domain. To test this, we checked the expression pattern of both genes in the adult head by inserting EGFP under the endogenous regulatory elements of DptB and GNBP-like3. Western blotting of 4-7 days old adult head extract showed EGFP expression from both genes, albeit the expression from GNBP-like3 locus were significantly lower than that from the DptB locus (**S8 Fig**). Immunostaining of DptB-EGFP genomic flies for EGFP and the neuronal gene bruchpilot (nc82 staining) revealed that EGFP expression is confined to the outer layer of the head, outside the central brain, where the head fat body is located (**Fig 5A** & **S8 Fig**). However, immunostaining for GNBP-like3 was not successful likely due to its very low expression level (data not shown). Therefore, we performed RNAscope, an *in* situ-based technique that uses several amplification steps, to detect GNBP-like3 mRNA expression in wild type and in *gnbp-like3−/−* as control. A specific signal for GNBP-like 3 was detected in the central brain (**Fig 5B**). Since mRNA expression does not necessarily mean protein expression, and within the central brain there are different cell types in addition to neurons, we further sought to verify the likely source of GNBP-like3 protein in the brain. To this end we inserted a 3HA-tag in the C-terminal end of GNBP-like3 endogenous locus using Crispr-Cas9 mediated-homologous recombination. Subsequently, using a multistep fractionation protocol previously developed, we isolated synaptosomes from adult fly head (**Fig 5C**). Western blotting showed HA-immunoreactive polypeptides in the purified synaptosomes (**Fig 5C**). To rule the possibility that HA-tagging had not altered expression or localization of the peptide, we also purified synaptic membrane and synaptic soluble proteins from wild type adult fly heads and performed proteomic analysis (**S9 Fig**). In proteomics, proteins that were detected at least 2 out of 3 independent purification were considered for further analysis (S5 Table). Approximately 105 proteins were detected specifically in the synaptic membrane fraction and among 43 known immune-related peptides, only a peptide from GNBP-like3, KVNEEMDDLSDQTWAADVVSSRN, was detected in the same fraction (**S9 Fig**). Taken together, these results suggest that consistent with their functional requirement, DptB is expressed in the head adult fat body, while GNBP-like3 is expressed in the neurons and likely present in the synaptic compartment. However, our analysis does not rule out that these peptides are not expressed at low levels in other head tissues.

**Figure 5.**
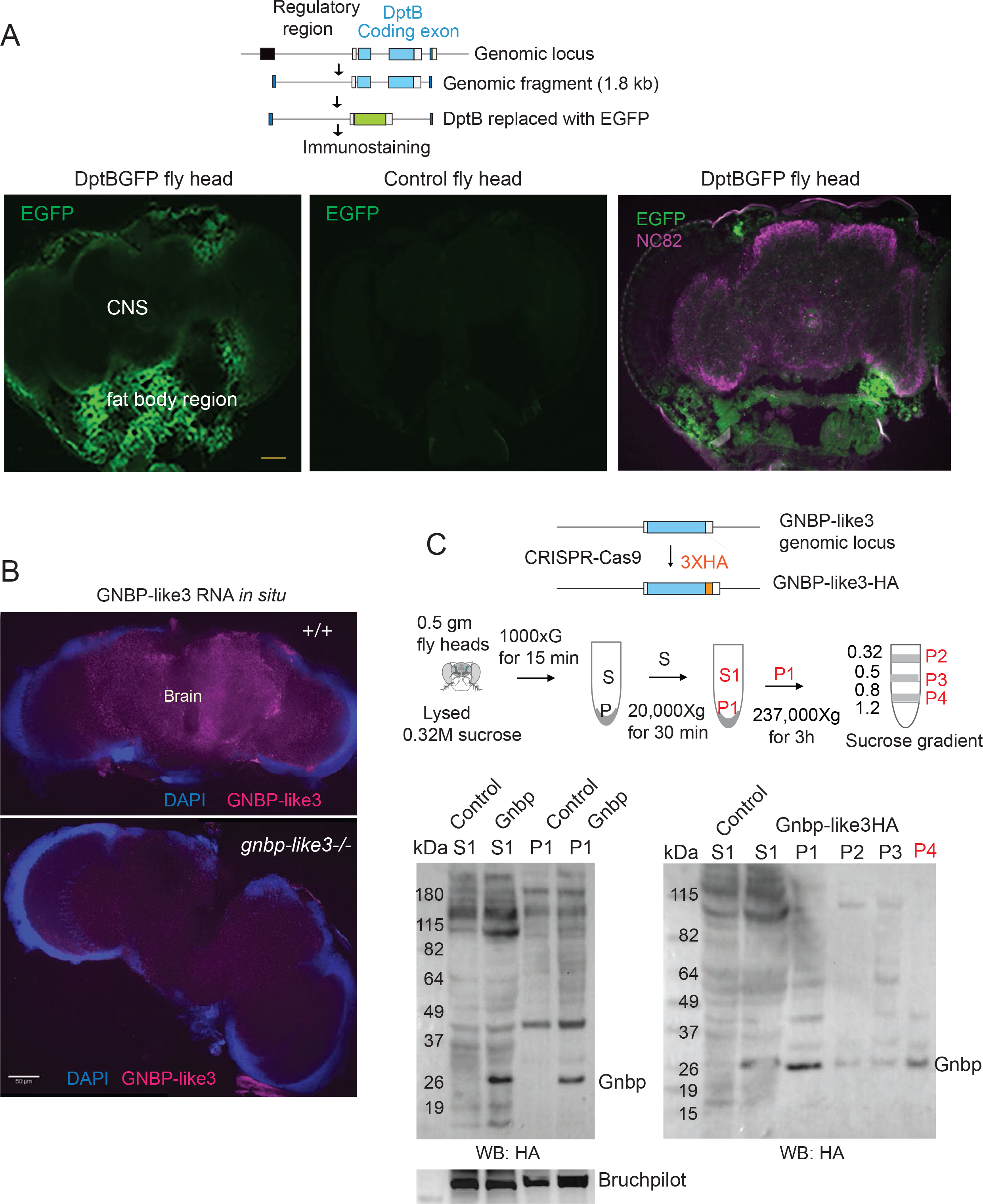
DptB is enriched in the head fat body and GNBP-like 3 in neurons. (**A**) EGFP expression when driven under DptB-regulatory sequences. Immunostaining reveals EGFP expression is confined to the outer layer of the head, outside the central brain (marked by nc82 antibody), where the head fat body is located. Wildtype brain serves as a control for non-specific immunoreactivity of anti-EGFP antibodies. (**B**) GNBP-like3 mRNA expression determined by RNAscope *in situ* hybridization shows expression in the central brain. Scale bar: 50 μm. (**C**) GNBP-like3 is present in synaptosome. Top. The HA-tag was introduced into the C-terminal end of GNBP-like3 by homologous recombination using Crispr-Cas9. Middle. Schematic depiction of synaptosome purification from adult fly heads. The P4 fraction, between 0.8M and 1.2M sucrose, is most enriched for synaptic protein. Bottom. Different fractions from the synaptosome purification were blotted for GNBP-like3. Wild type flies serve as a control for HA-antibody specificity.

### DptB and GNBP-like3 attenuate bacterial growth

*Drosophila* genome encodes many peptides that are upregulated upon bacterial infection, however, they may or may not have antimicrobial activity. DptB is a 120aa long-peptide with 52% similarity to the Gly-rich domain of antimicrobial peptide DptA and 37% similarity to the second Gly-domain of the Attacin family antimicrobial peptide AttacinA. GNBP-like3 shares homology to pattern recognition receptors that bind to components of bacterial cell wall and activates innate immune response [22,23,24]. However, GNBP-like3 is distinct from other GNBPs in several ways. First, most GNBPs are longer than 400 aa, while GNBP-like3 is only 152 aa long and lacks the C-terminal sequence present in most GNBPs. Second, unlike canonical GNBPs, whose expression level is unaltered, GNBP-like3 is upregulated following bacterial infection, a feature of antimicrobial peptides.

Although assumed, however, to our knowledge, there is no report of direct antimicrobial activity of either DptB or GNBP-like3. Therefore, we set out to compare the effect of GNBP1, GNBP-like3 and DptB on bacterial growth, with that of a well characterized AMP, Drosocin. To accomplish this, we used an inducible bacterial expression system where the AMPs or the control mCherry were placed under L-arabinose inducible pBAD promoter (**Fig 6A**). However, when grown in synthetic media containing L-arabinose that induces expression of GNBP-like3 or DptB, bacterial growth was significantly reduced, like that of Drosocin (OD600 after 22h: mCherry, 0.78±0.006; GNBP-like3, 0.48±0.005; DptB, 0.47±0.003; and Drosocin 0.48±0.002). Surprisingly, bacteria harboring GNBP1 did not grow at all in synthetic media containing L-arabinose (OD600 after 22h 0.11± 0.002), indicating although GNBP1 and GNBP-like3 share homology, they may be functionally distinct, and GNBP-like3 and DptB may act as bacteriostats.

**Figure 6.**
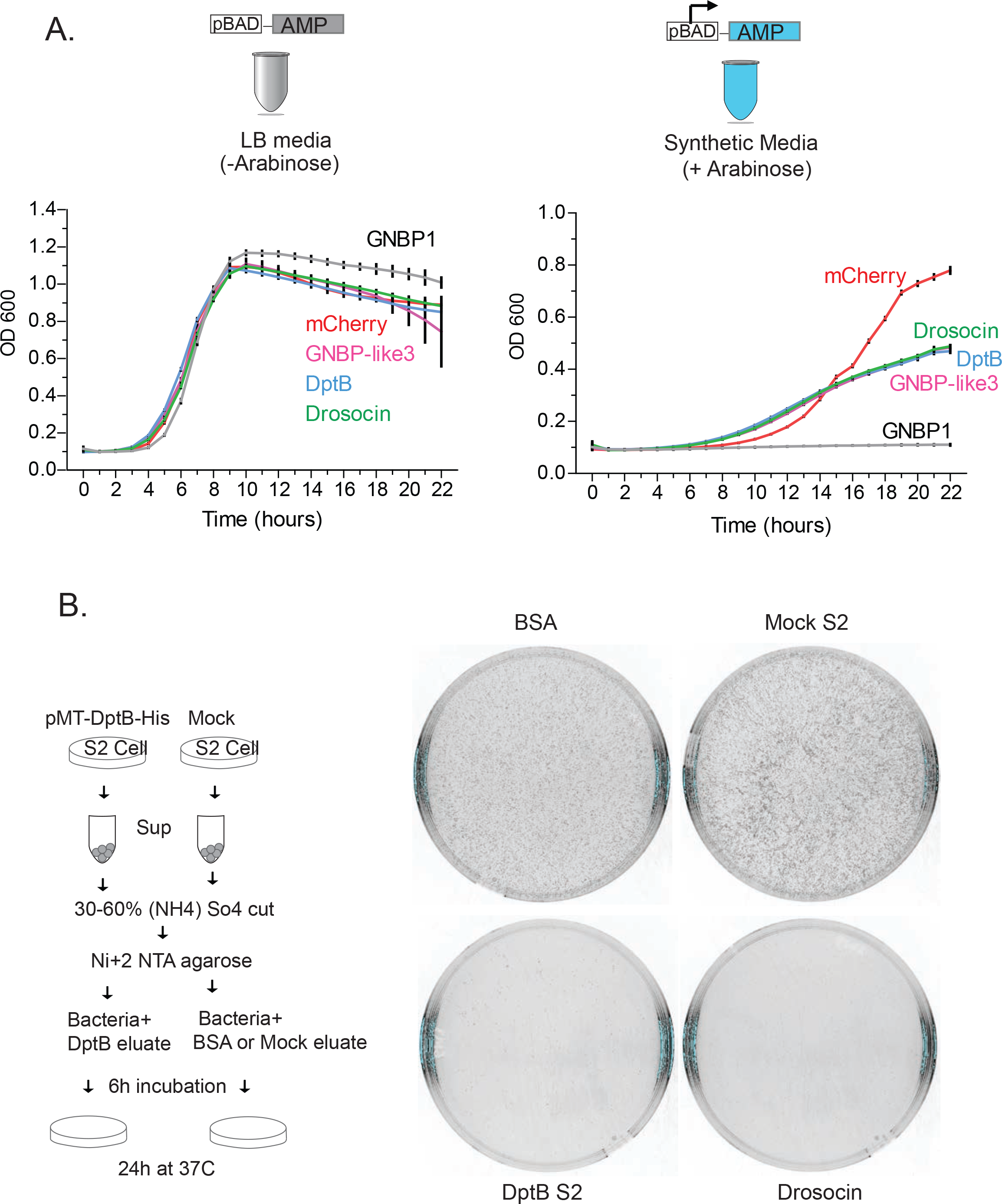
DptB and GNBP-like3 have antimicrobial activity. (**A**). Growth of bacteria bearing GNBP-like3, DptB, Drosocin, GNBP1, and mCherry as a control, in two different culture conditions: the rich LB media (left), and synthetic media with L-arabinose to induce the expression of the constructs (right). Bacterial growth is represented by OD600 over time. (**B**). Left, schematic representation of the experimental design. Right, bacterial plates showing that the fraction containing DptB slows bacterial growth. The known antimicrobial peptide Drosocin is used as a positive control. DptB: DiptericinB, GNBP1: Gram-Negative Binding Protein 1, GNBP-like3: Gram-Negative Binding Protein like3.

To assess more directly the effect of DptB and GNBP-like3 on bacterial growth, we attempted to purify them from S2 cells expressing C-terminal HA-tagged DptB and GNBP-like3. The DptB-HA-tagged protein could be detected in the total cell lysate and in the media upon ammonium sulfate precipitation (**S10 Fig**). Interestingly, as in the brain, the GNBP-like3 had very low expression in S2 cells and could only be detected in the total cell lysate (**S10 Fig**), but not in the media, and its low expression level prevented further analysis. To determine whether the secreted DptB peptide possess any biological activity, we expressed a C-terminal histidine-tagged version and enriched for the secreted peptide from the media by in Ni+2 affinity column (**Fig 6B**). Same amount of untransfected S2-cell media were similarly purified as negative control and the synthetic antimicrobial peptide Drosocin was used as a positive control. Incubation of the Ni+2 affinity bound fraction from DptB expressing cells resulted in inhibition of bacterial growth compared to the untransfected or BSA control, suggesting that the secreted fragment of DptB indeed possess anti-bacterial growth inhibitory activity (**Fig 6B**). Like direct bacterial expression, addition of DptB to bacteria attenuated growth, but did not abolish it. Taken together these results suggest that DptB and GNBP-like3 have bacteriostatic activity.

## Discussion

### Normal function of antimicrobial peptides in the adult brain

We set out to identify genes that are specifically involved in long-term memory. Our experimental set up expected to identify immediate early genes that are transcriptionally induced by behavioral training that produces long-term memory. We observed that some of the known memory related genes that were changed in the trained groups (**S3 Fig**), were also changed upon starvation or exposure to sorbose that does not produce robust long-term memory. This suggests that genes that are changed in some of the control conditions may also be important for memory, although they fail to satisfy the stringent criteria we have set up. Moreover, our analysis would miss genes that are involved in memory, but not transcriptionally induced, as illustrated by GNBP-like3 and other memory related genes (**S3 Fig**). Remarkably, despite the stringency of our analysis, a few immune peptides are consistently up regulated in groups that are trained to form stable long-term memory.

There is increasing evidence that components of the immune system also function in the nervous system [1,2,3,4]. Similarly, repurposing of AMPs, although rare, is not unprecedented. For example, mammalian β-defensin acts as a ligand for the melanocortin receptor 1 (Mc1r) to control melanin synthesis [25]. In *Drosophila*, AMPs, such as Metchnikowin (Mtk), Drosocin and Attacin, are implicated in regulation of sleep [5]; moreover, the innate immune receptor PGRP-LC is involved in homeostatic plasticity of neuromuscular junction synapse [2]. However, to our knowledge, this is the first time that AMPs made in different tissues in adult head have been found to be involved in modulating long-term associative memories. More recently, Dpt, a different antimicrobial peptide, has been shown to be important for a form of nonassociative learning, where ethanol preference is modified upon exposure to predatory wasp [6].

### Why immune peptides?

For most animals, including insects such as *Drosophila melanogaster*, the ability to remember a potential food source or modulate reproductive behavior based on prior experiences is a valuable trait. Both feeding, and copulation expose the inside of the animal to the external environment. Therefore, these events are likely to engage the immune system in preparation for the exposure to external agents, including pathogens. We postulate that DptB, GNBP-like3, and other AMPs are upregulated in the body to deal with immune challenges. Subsequently, over evolutionary time, in addition to their protective roles in immunity, some immune related genes were repurposed to act as modulators of nervous system function. The nervous system perhaps co-opted these immune genes to convey and store information about specific aspects of experiences. The type of information represented by these peptide signals remains unclear at this stage.

Interestingly, we find that while both DptB and GNBP-like3 have similar requirement for long-term courtship suppression memory, their requirement in associative appetitive memory is different. What accounts for this differential dependency on a set of molecules? It is possible that the animals prioritize survival over reproductive success, and therefore remembering a food source involves several molecules that can compensate for the absence of each other. This is consistent with the fact that appetitive memory is quite robust and requires only one training for 5 minutes, while to elicit long-term courtship suppression memory requires multiple training lasting for 6 hours. In any event, these observations raise the possibility that in addition to common molecular processes, different types of memories may have unique molecular requirements. Indeed, a different group of immune peptides are up-regulated when *Drosophila* forms memory of a predator, such as wasp [6].

A key unanswered question of considerable interest is how and where DptB, and GNBP-like3, act to influence memory? How these AMPs act at molecular and cellular level to modulate memory is key to understand in greater depth the relationship between AMPs and memory. An important step in this direction would be to identify the “receptors” of these AMPs in the adult nervous system. By receptor we mean either cell surface or intracellular components that transforms the AMP activity to a neuronal (AMPs may also act upon non-neuronal cell) response. Identification of the “receptor” would uncover in which cell population these AMPs act, how they change cellular function and when the AMP-mediated modulation of the cellular function is important for memory.

### Additional significance of immune peptides in the brain

Is there additional significance to the observation that AMPs modulate nervous system functions? Curiously, some neuropeptides, like NPY, possess antimicrobial activity, and innate immunity-related peptides are expressed in the mammalian brain [26]. However, the expression of AMPs in the brain is often associated with dysfunction. For example, overexpression of antimicrobial peptides in *Drosophila* brain accelerates neurodegeneration [27]. Recently, Aβ-42, the truncated product of amyloid-precursor-protein (APP) and a causative agent for Alzheimer’s disease, has been postulated to be an AMP [28]. In this view, although the central nervous system is isolated by the blood-brain-barrier, these AMPs are present in the brain to fight invading pathogens, or the AMPs are produced in the brain in response to inflammation or other stress. We speculate that AMPs are made in the brain, not necessarily exclusively for immune related functions, but to act as canonical neuromodulators to regulate nervous system functions. Indeed, the requirement of GNBP-like3 in neurons and its presence in synaptosomes are consistent with such a possibility. That some AMP expression eventually leads to dysfunction is perhaps an unintended consequence of a normal process [29].

## Methods

### Fly strains

Gal4 lines used in behavioral experiments: S32, head fat body driver (BDSC #8527); S106, body fat body driver (BDSC #8151); 3.1Lsp2, head and body fat body driver is a gift from Dr. Brigitte Dauwalder; Actin-geneSwitch (255B) was a gift from Dr. John Tower; for glial cell driver we used RepoGal4 (BDSC #7415); for neuronal drivers we used ElavGal4 (BDSC #458) and Elav-geneSwitch which is a gift from Haig Keshishian. UAS RNAi lines: mCherry (BDSC #35785), DptB (BDSC #28975), luciferase (BDSC #35788), Dpt (BDSC #53923), AttA (BDSC #56904), AttB (BDSC #57392), AttC (VDRC #101213), Drs (BDSC #55391), Def (BDSC #29524), Mtk (BDSC #28546), Dro (VDRC #105251), GNBP-like3 (VDRC #107358). Flies were maintained at 25 °C, with a 12:12 light:dark period.

### Isolation of single fly head RNA

After indicated behavioral treatment, flies were flash frozen in liquid nitrogen. Single animals were then isolated on dry ice to avoid thawing and collected in 1.5 mL Eppendorf tubes, which were immediately replaced in liquid nitrogen. To separate heads from bodies the flies were either vortexed or individually cut off in a dry ice block. RNA was then collected with the Maxwell^®^ 16 LEV simplyRNA Tissue Kit (Promega). Briefly, the given tissue was homogenized in 50 μl of Homogenization Buffer for 15 seconds using an RNAse/DNAse free pestle (VWR) and electric pestle mixer (VWR). An additional 150 μl of Homogenization Buffer was then added followed by addition of 200 μl of Lysis Buffer. Samples were then vortexed for 15 seconds and spun down. The provided cartridges were used according to the protocol to purify the RNA in 30 μl of nuclease free water. After completion of the RNA isolation, samples were placed in a vacuum centrifuge for approximately 8 minutes to achieve a volume of 5-7 μl.

### mRNA-Seq

Samples were all collected using RNA isolation as described above on single female fly heads. In appetitive memory paradigm for each condition the number of samples in the analysis were as follow: Untreated: 5 individual heads; starved: 5 individual heads; 1 hour post training (Sucrose): 6 individual heads; 4 hours post training (Sucrose): 5 individual heads; 4 hours post training (Sucrose II): 6 individual heads; 4 hours post training (Sorbose): 6 individual heads. In male courtship suppression paradigm 6 individual virgin male fly heads were collected after the following conditions: 1 hour after 1X mock training, 1 hour after 1X training with an unreceptive female; 1 hour after 3X mock training, and 1 hour after 3X training with an unreceptive female. Library preparation and submission was conducted by the Molecular Biology Core at the Stowers Institute. For analysis of the data we used a protocol described by Trapnell et al. [30]. Briefly, RNA-seq analysis was done using TopHat v1.4.1 [30] and Bowtie v0.12.7. Only uniquely mapping reads to fly genome UCSC dm3 were used. Fly transcript annotations were from Ensembl 65. Differentially expressed genes were called with an adjusted p value (FDR) < 0.05 by cuffdiff v1.3.0.

### Quantitative PCR

The qPCR was performed as described in Gill et.al [31] except random hexamer was used for cDNA synthesis. The following primers were used for DptB qPCR: “ACTGGCATATGCTCCCAATTT” and “TCAGATCGAATCCTTGCTTTGG”. The following housekeeping genes were used: GAPDH (“AGGGAGCCACCTATGACGAAATCA” and “AGACGAATGGGTGTCGCTGAAGAA”), RPL32 (“AGCGCACCAAGCACTTCATC” and “GACGCACTCTGTTGTCGATACC”), and TBP (“TCCAGACTGGCAGCGAGAAAGTAT” and “AACTTGACATCGCAGGAGCCG”).

### RT-PCR

Total mRNA was isolated using Trizol™ and cDNA was synthesized using the first-strand cDNA synthesis kit (Invitrogen). PCR was performed using the following primers: DptB (“TGGGCTTATCCCTATCCTGA” and “ATAGGGTCCACCAAGGTGCT”), AttA (“ACAATGTGGTGGGTCAGGTT” and “GTCAAGGAAGCACCATGTCC”), AttB (“ATTGGCAATCCCAACCATAA” and “TGTCCGTTGATGTGGGAGTA”), AttC (“GGTATCCACTCCCGTAACCA” and “TCCAATTGCTTACCCAATCC”), and Actin42A (“TGGACAGGTCATCACCATCGGAAA” and “TTGTAGGTGGTCTCGTGAATGCCA”).

### Immunohistochemistry

Flies were decapitated in CO_2_. Fly heads were frozen in OCT. 25 p,m frozen sections of these fly heads were fixed in 4% paraformaldehyde in TBS-Tri (0.05% Triton-X-100) for 10 min at room temperature. After washes in TBS-Tween (0.05% Tween20), the sections were blocked in TBS-Tween-NGS (Normal Goat Serum, 10%) at room temperature for 1 h, and then incubated with primary antibody in TBS-Tween-NGS overnight at 4°C. The second day, sections were washed in TBS-Tween and then incubated in secondary antibody in TBS-Tween-NGS overnight at 4°C. After additional washes, sections were mounted in Vectashield medium with DAPI and coverslips. Rabbit-anti-GFP (MBL #598), mouse-anti-bruchpilot (DSHB nc82) were used as primary antibodies. Alexa-594 coupled antirabbit, and Alexa-647 coupled anti-mouse secondary antibodies were used. Confocal images were acquired using Ultraview Spinning Disc confocal microscope.

### Generation of DptB and GNBP-like3 knock-out flies

DptB gRNA (target sequence “GCAGATGAGATTCCCCAATTGGG” and “GGCGGTGGAAGCCCAAAGCAAGG”) and GNBP-like3 gRNA (target sequence “CGCTGAAGCCTCAGTTCTCG” and “GACGCTGGACTTATCGCAAM”) were cloned into vector pU6.2, which were then injected into the attP2 line (BDSC #25710), or nos-Cas9 (III-attP2) line by BestGene, respectively. In case of DptB gRNA flies were crossed to nos-cas9 flies to obtain founders. KO were screened and confirmed by PCR and sequencing. Then KO flies were backcrossed to Canton-S background.

### Generation of DptB and GNBP-like3 eGFP flies

DptB promoter-eGFP genomic construct was made, by Gibson assembling these fragments: (1) linearized pattB vector; (2) DptB upstream (intergenic region between Dpt and DptB, presumably contains promoter and region for DptB); (3) eGFP; (4) DptB downstream (intergenic region between DptB and CG43109). This construct was injected into attP2 line (BDSC #25710) to generate DptB promoter-eGFP line. GNBP-like3 promoter-eGFP genomic construct was made, by Gibson assembling these fragments: (1) linearized PHD-dsRed vector; (2) GNBP-like3 upstream (~1.2 Kb, presumably contains promoter for GNBP-like3); (3) eGFP; (4) GNBP-like3 downstream (~1.2 Kb). This construct, together with gRNAs were injected into nos-Cas9 (III-attP2) line by BestGene to generate GNBP-like3 endogenous locus substitution by eGFP.

### Courtship behavior

Male virgin flies were taken soon after eclosion and were kept in isolation for 5 days. In the fifth day, each male was introduced in a chamber with a 5-day-old virgin female and recorded them for 30 minutes. Manually, we studied the parameters described in Ejima and Griffith, 2007; Courtship latency: the time between introduction of the male fly and the first attempt to court, which is measured as orientation of the male fly towards the female; Copulation latency: time between pairing and first successful mounting; Copulation duration: total duration of the first mounting episode.

### Courtship Conditioning

The male courtship suppression conditioning was carried out as described previously [32]. Briefly, male flies were collected shortly after eclosion and housed individually for 5 days. For GeneSwitch experiments, flies were fed 1 mM RU-486 (mifepristone, SigmaM8046) in 2% sucrose for 3 days (starting with 2 days old flies) before training. Three spaced trainings were performed on day 5. Every training session lasted for 2 hours with 30 min intervals. During training, mated females were introduced to the males. For mock training male flies were introduced into the training tubes without any female flies. Flies were tested 15 minutes and 24 hours after training. The flies’ behaviors were recorded for 15 min by camera and analyzed by ImageJ.

### Sucrose preference

The two-choice tests were performed essentially as previously described in McGinnis et.al [10]. Briefly, 1 to 4-day-old flies were food-deprived in groups of 50 for 18-24 h in tubes containing kimwipes wet with 3 ml of water. 1% agarose (Sigma) was mixed into sugar solutions along with red or green food dye (1%, McCormick), and 15 μl drops were pipetted into 60-well minitrays (Thermo Scientific). 50 flies were allowed to feed for 5 min in complete darkness and the color in their abdomens was assessed under a dissecting microscope. Preference index was calculated as (number of flies eating sugar)/(total number of flies). To rule out the color bias half of the experiments had the colors reversed.

### Olfactory-Appetitive Conditioning

The appetitive associative training was carried out as described previously [32]. Briefly, ~100 flies were food deprived for 20-24 hours before conditioning. Flies were transferred to the –CS tube and exposed to an odor for 2 min. After 30 s of air stream, the flies were shifted to the +CS tube in the presence of the second odor for 2 min. Memory was tested 24 or 48 h after training. For 24-hour test, flies were given standard cornmeal food for 3 to 6 h after training and starved again for 17-20 h before testing. For 48-hour memory test, flies were given standard cornmeal food for 17–24 h after training and then were starved for 24–30 h prior to test. During testing, animals were given 2 min to choose between the two odors. Different group of flies were trained in a reciprocal experiment in which the −CS/+CS odor combination was reversed (3-Octanol or 4-Methylcyclohexanol). The memory index (MI) is calculated as the number of flies in the reward odor minus the number of flies in the control odor, divided by the total number of flies in the experiment. A single MI value is the average score of the first and the reciprocal experiment.

### Heat Box Paradigm

The heat box operant conditioning was performed using the apparatus and protocol of Wustmann et al. [15] and as described in Gill et. al [31].

### RNAscope *in situ* hybridization

The brains were dissected in PBS and fixed immediately in 4% PFA for 30 minutes. After washing with PBS-T (PBS + Triton 0.1%), the brains were dehydrated by sequentially washing (5 min/wash) in 25%, 50%, 75% and 100% methanol and eventually left in 100% MeOH at −20°C for 3 hours. Subsequently, the brains were incubated in protease III solution at room temperature for 20 minutes, washed in PBS-T and incubated with the probes at 50°C overnight. All subsequent washes were done using 0.2X SSCT for 5 minutes 3 times. The brains were washed to remove the unbound probe, fixed with 4% PFA for 10 minutes. The brains were then washed and incubated with Amplifier1 at 42°C for 30 minutes, Amplifier2 at 42°C for 15 minutes, Amplifier3 at 42°C for 30 minutes and Amplifier4AltA at 42°C for 15 minutes. Between each amplification step the brains were washed. After the final wash, the brains were mounted in Vectashield mounting media with DAPI. Images were acquired in an Ultraview Spinning Disc confocal microscope and the images were processed using ImageJ. The experiment was repeated more than 3 times with independent biological samples.

### Synaptosome from adult fly head

Synaptosomes were purified as described in Majumdar et.al [8]. Briefly all Sucrose solutions were made in Buffer A [10mM Tris.HCl (pH7.5)]. Adult fly heads were crushed in 0.32 M sucrose buffer (2ml/0.5gm of head) and centrifuged twice at 1000Xg for 20 min to separate the nucleus and other heavier cellular components from the membrane and soluble proteins. The supernatant (T) was centrifuged at 15000Xg for 15 min and the resulting pellet was resuspended and centrifuged again 15000Xg for 10 min to obtain washed crude synaptosome fraction (P1). The P1 fraction was resuspended in 0.32 M Sucrose buffer and 1 ml of the resuspended pellet was loaded on top of a 9.9 ml 0.5 M, 0.8 M and 1.2 M sucrose buffer step gradient and centrifuged at 100,000Xg for 3 hours in a Beckman SW40 rotor. The interface between 0.8 and 1.2 M sucrose was collected, pelleted, resuspended in 1 ml of 0.32 M sucrose solutions and again loaded. The interface between 0.8 and 1.2 M sucrose was collected, diluted to 8 ml with 0.32 M Sucrose, loaded on top of 4 ml 0.8 M sucrose buffer and centrifuged at 230,000xg for 22 min in SW40 rotor to obtain purified synaptosome (P2). The pellet was extracted with 1% NP40 and 1% Triton X-100, 10 mM Tris.HCl (pH7.5) buffer. The resuspended pellet was centrifuged 15,000xg for 15 min and the supernatant was used as soluble synaptic fraction. The pellet containing purified synaptic membrane was extracted with buffer containing 1%NP-40, 1%TritonX-100 and 1%SDS, 10mM Tris.HCl (pH7.5) (P3).

### Antimicrobial activity

The open reading frame of DptB, GNBP-like3, Drosocin and GNBP1 were cloned into bacterial pDiGc expression vector [33]. NheI and AleI restriction sites were introduced into the 5 and 3’ end of the cDNA and introduced into the same sites in pDiGc vector. To study bacterial growth, bacteria carrying the pDiGc-AMPs were grown in LB (pH=7) or MgM-MES media (pH=5) in presence of ampicillin overnight at 37°C. The cultures from log phase were diluted to 0.1 OD_600_ either in LB media or in synthetic media containing L-arabinose (0.2% w/v, Sigma). The cultures were grown in 96-well plates and using a spectrophotometer OD_600_ was measured continuously for 22 hours at 37°C. The absolute value of ODs after every hour was plotted mean +/− Standard Deviation of the mean.

To purify His-tagged DptB, C-terminal 6XHis tagged DptB was cloned placed Cu+2 inducible metalothionin promoter and the construct was inserted in S2 cell genome to make stable cell line. 500 ml of S2 cell in SPX/PS media (a protein free media) was induced with 500 uM CuSO4 and after 48 h cells were harvested at 5000 rpm, 15 min at 4°C. The supernatant was subjected to 0-30% and 30-60% ammonium sulphate cut. The pellet of 30-60% cut was dissolved in 10 ml PBS and dialyzed against 1 lit PBS at 4°C overnight. The dialyzed material was incubated with 3 ml Ni+2-NTA agarose, washed and eluted with 300 mM imidazole and dialyzed against sterile 0.9% NaCl.

To assess anti-microbial activity, bacteria was grown to log phase (OD_600_ 0.5-1.0), collected by centrifugation at 8000 rpm for 30 s, washed with 0.9% NaCl and resuspended with 0.9% NaCl to 0.002A. 15 μl of bacteria was added to 15 μl protein solution, incubated at room temperature for 5 h and spread into a LB-plate. Different dilutions were spread to ensure easy visualization of the colonies, and same dilutions were compared to measure the effect on growth, if any.

### Statistical analysis

All statistical analysis was performed using Graphpad Prism 5. All data met the assumption of homogeneity of variance, therefore unpaired two-tailed t-test or one-way analysis of variance (ANOVA) was performed, with Bonferroni post-hoc test between pairs of samples. ANOVA tests for significance were performed a probability value of 0.05 and more stringent values are listed in each figure where applicable. For all experiments, each *n* is considered a biological replicate; separate trials used independent samples of genetically identical flies.

### Processing of Genomic Data Sets

1. Courtship suppression microarray 0 h & 24 h [34](Kitamoto et al., 2006)
2. Courtship suppression mRNA seq 24 h (Winbush et al., 2012)[35]
3. Appetitive Sucrose or L-sorbose 4 h after training.
4. Appetitive Sucrose 1 h & 4 h after training.
5. Courtship suppression 1X and 3x training

The genomic data was processed using tools developed by Benjamini and Hochberg (1995) [36], and Robinson et al. (2010) [37]. The courtship suppression microarray data set representing studies of Kitamoto et al. 2006 was found under ArrayExpress accession number: E-GEOD-4032 and subsequently retrieved from GEO using accession numberGSE4032. The data was analyzed with GEO2R based on descriptions of the twelve samples on the GEO Accession viewer. They are as follows:

| id | tissue | name | genotype | time | conditioning | replicate | description |
| --- | --- | --- | --- | --- | --- | --- | --- |
| GSM92389 | whole head | extract#5_le1 | OR | 0 | 0 | 1 | 0hr after non-conditioning |
| GSM92390 | whole head | extract#6_le1 | OR | 0 | 0 | 2 | 0hr after non-conditioning |
| GSM92395 | whole head | Sample11_e1_le1 | OR | 0 | 0 | 3 | 0hr after non-conditioning |
| GSM92385 | whole head | extract#1_le1 | OR | 0 | 1 | 1 | 0hr after conditioning |
| GSM92386 | whole head | extract#2_le1 | OR | 0 | 1 | 2 | 0hr after conditioning |
| GSM92393 | whole head | Sample9_e1_le1 | OR | 0 | 1 | 3 | 0hr after conditioning |
| GSM92391 | whole head | extract#7_le1 | OR | 24 | 0 | 1 | 24hr after non-conditioning |
| GSM92392 | whole head | sample#8_le1 | OR | 24 | 0 | 2 | 24hr after non-conditioning |
| GSM92396 | whole head | Sample12_e1_le1 | OR | 24 | 0 | 3 | 24hr after non-conditioning |
| GSM92387 | whole head | extract#3_le1 | OR | 24 | 1 | 1 | 24hr after conditioning |
| GSM92388 | whole head | extract#4_le1 | OR | 24 | 1 | 2 | 24hr after conditioning |
| GSM92394 | whole head | Sample10_e1_le1 | OR | 24 | 1 | 3 | 24hr after conditioning |

Data was analyzed using GEO2R under default settings. Data was exported with log transformation and the p-values were adjusted with Benjamini & Hochberg (1995)[36]. Probes with a p-value less than 0.05 were considered significant. Those probes were translated to genes using the NetAffx Analysis Center provided by AffyMetrix NetAffx. Some probes translated into multiple genes; in that case, each gene was assigned the p-value corresponding to the probe. The reverse also occurred: sometimes multiple probe sets would target the same single gene, thus the gene may have multiple p-values assigned to it. In such cases, the lowest p-value was assigned to highlight the potential significance of that gene. For the clear majority of genes, neither case occurred. Most probes corresponded to one gene and one value.

Data from Winbush et al. (2012) [35], representing a courtship conditioning paradigm was downloaded from SRA under accession number SRA055890. Data was aligned to dm6 using tophat2 and Ensembl 84 gene models. The edgeR library (Robinson et al., 2010)[37] in the R environment was used for normalization and to assess differentially expressed genes between conditions. P-values were adjusted for multiple hypothesis testing the method of Benjamini and Hochberg (1995).

Data sets 3, 4, and 5, representing Appetitive conditioning with either Sucrose or L-sorbose (3) or a replicate experiment with Sucrose at 1 and 4 hours (4), or a Courtship conditioning paradigm (5) were processed identically as described for the data from Winbush et al. (2012)[14] as described above.

## Acknowledgments

KS is funded by the Stowers Institute for Medical Research. The project was conceived by KS. RBA and JW performed all the experiments. JG performed the mRNA sequencing. RW and CS performed the sequence analysis. The paper was written by KS and RBA. We are grateful to the following individuals for their help in various aspects of the work: Sean McKinney for writing the code for automatic behavioral analysis; Brian Slaughter, Jay Unruh and Yunming Yu for help with RNA *in situ*; Jose Lopez Hernandez (Fibo) for help with sequence analysis; Anita Saraf for proteomic analysis; Kyle Patton from Si lab for his comments on the manuscript.

---

**Supplementary Table S1. Genes that are up- or down-regulated after 1X or 3X training in male courtship suppression paradigm.** In red, genes up-regulated; in green, down-regulated. Antimicrobial peptides are highlighted in bold.

**Supplementary Table S2. Genes up- or down-regulated after different appetitive associative conditionings.** In red, genes up-regulated; in green, down-regulated. Antimicrobial peptides are highlighted in bold.

**Supplementary Table S3. Genes up- or down-regulated after different male courtship suppression trainings, and different mRNA sequencing methods.** Antimicrobial peptides are highlighted in bold.

**Supplementary Table S4. Genes that are significantly changed under various behavioral conditions.** Significant change (up or down) in mRNA level compared to control is indicated with √. The change in mRNA level after training in courtship paradigm (0h and 24h, microarray, and RNAseq) from other groups, and 1h after training in appetitive associative memory paradigms are also included in the table. Please see processing of mRNA sequencing data in the methods section for detail analysis. Antimicrobial peptides are highlighted in bold.

**Supplementary Table S5. The mass-spec analysis of purified synaptic membrane and synaptic soluble fractions.** Proteins that were detected at least two samples are included in the list. Mb stands for synaptic membrane fraction and sol stands for soluble fraction. Please see the synaptosome from adult fly head in methods section for detail information.

**Supplementary Figure S1.**
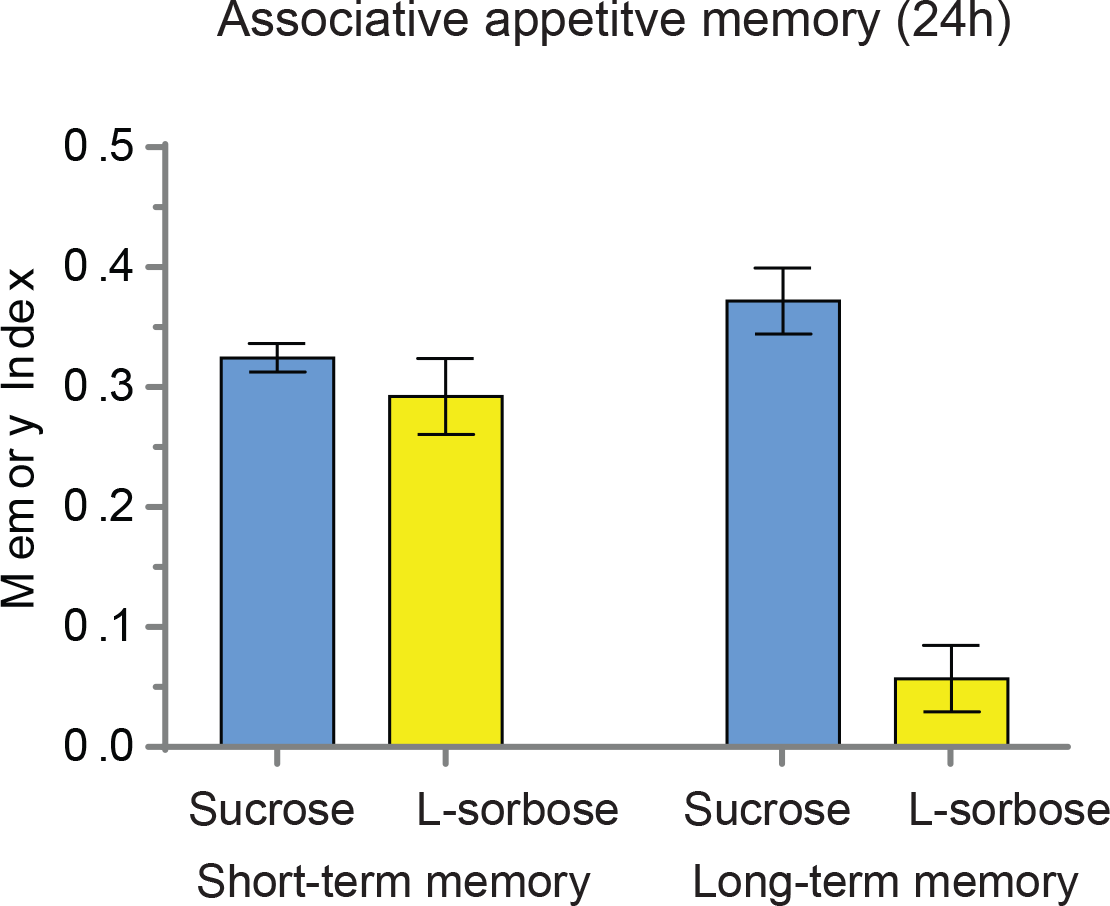
Sucrose produces stronger long-term memory than L-sorbose in associate appetitive paradigm. Memory index of flies trained with sucrose, and L-sorbose. The data are plotted as mean ± SEM.

**Supplementary Figure S2.**
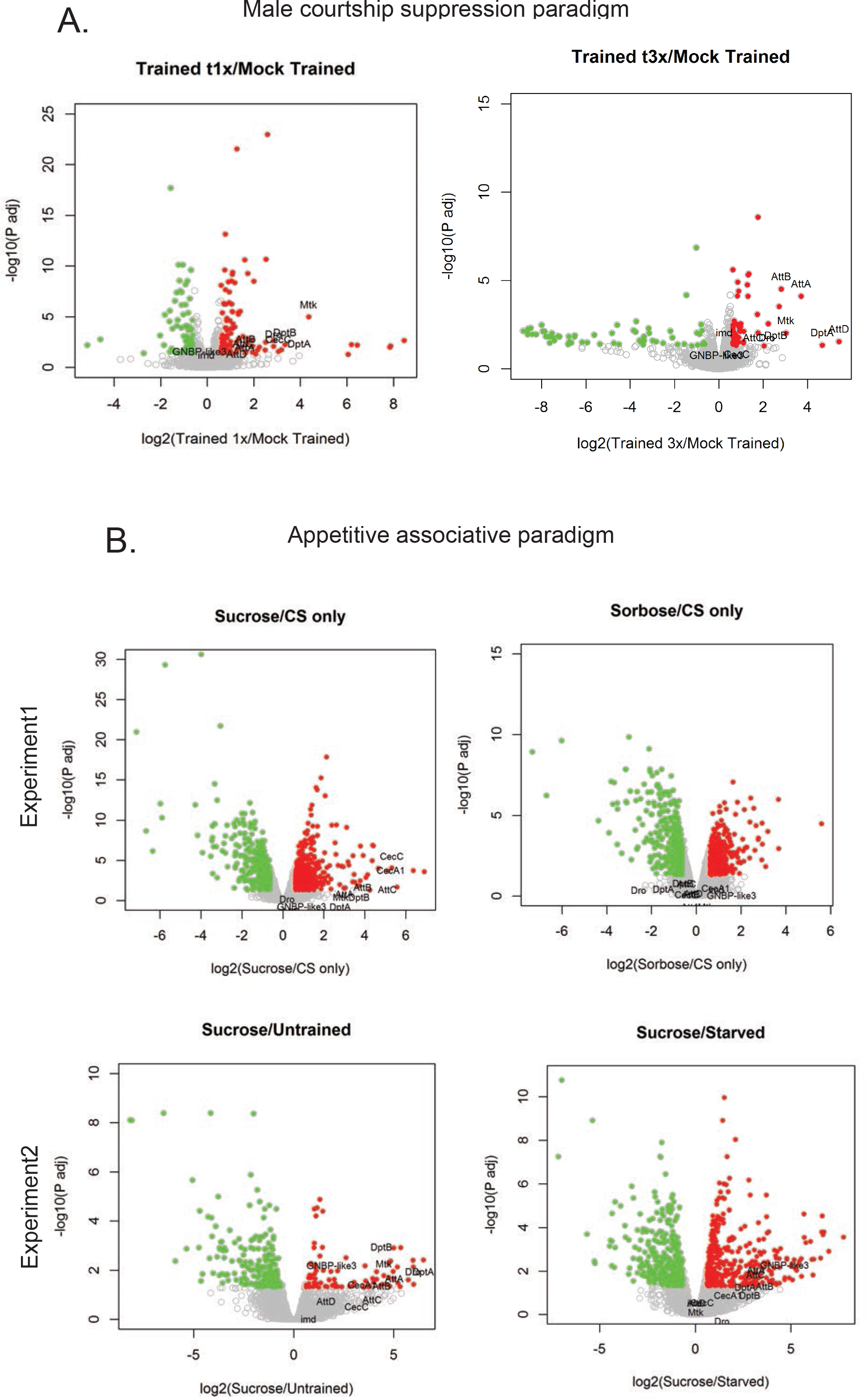
Gene expression changes under different training conditions. (**A**) Volcano plots showing genes up- or down-regulated after 1X training (left), or 3X training (right), compared to mock trained control. (**B.**) Volcano plots showing genes up- or down-regulated after sucrose or sorbose training compared to the CS only control in the experiment 1(top); and genes up- or down-regulated after sucrose training compared to the untrained or starvation controls in the experiment 2 (bottom).

**Supplementary Figure S3.**
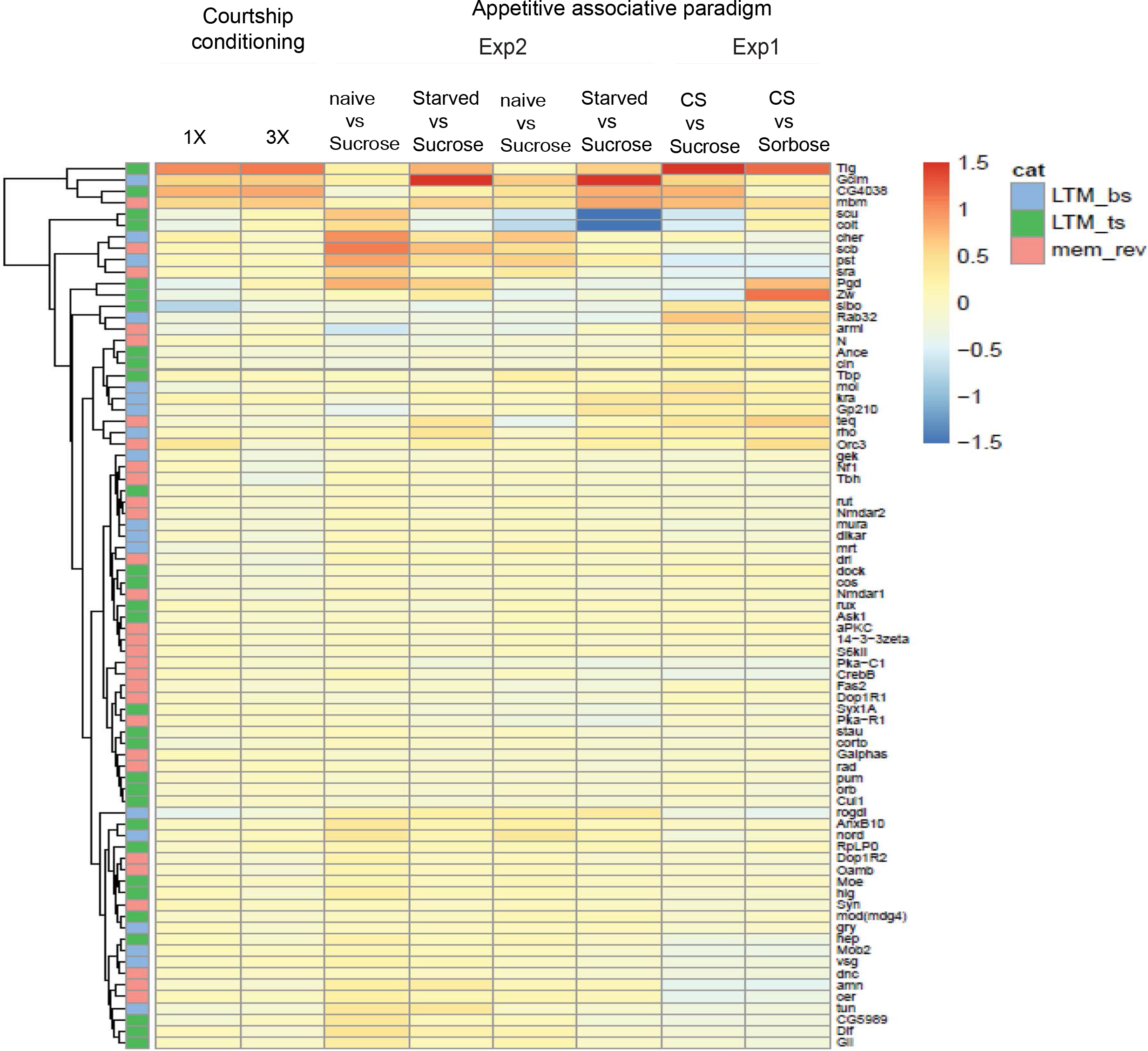
Expression of know memory genes under different training conditions. Heat map of the expression of known memory genes measured 1 hour or 4 hours after different training conditions.

**Supplementary Figure S4.**
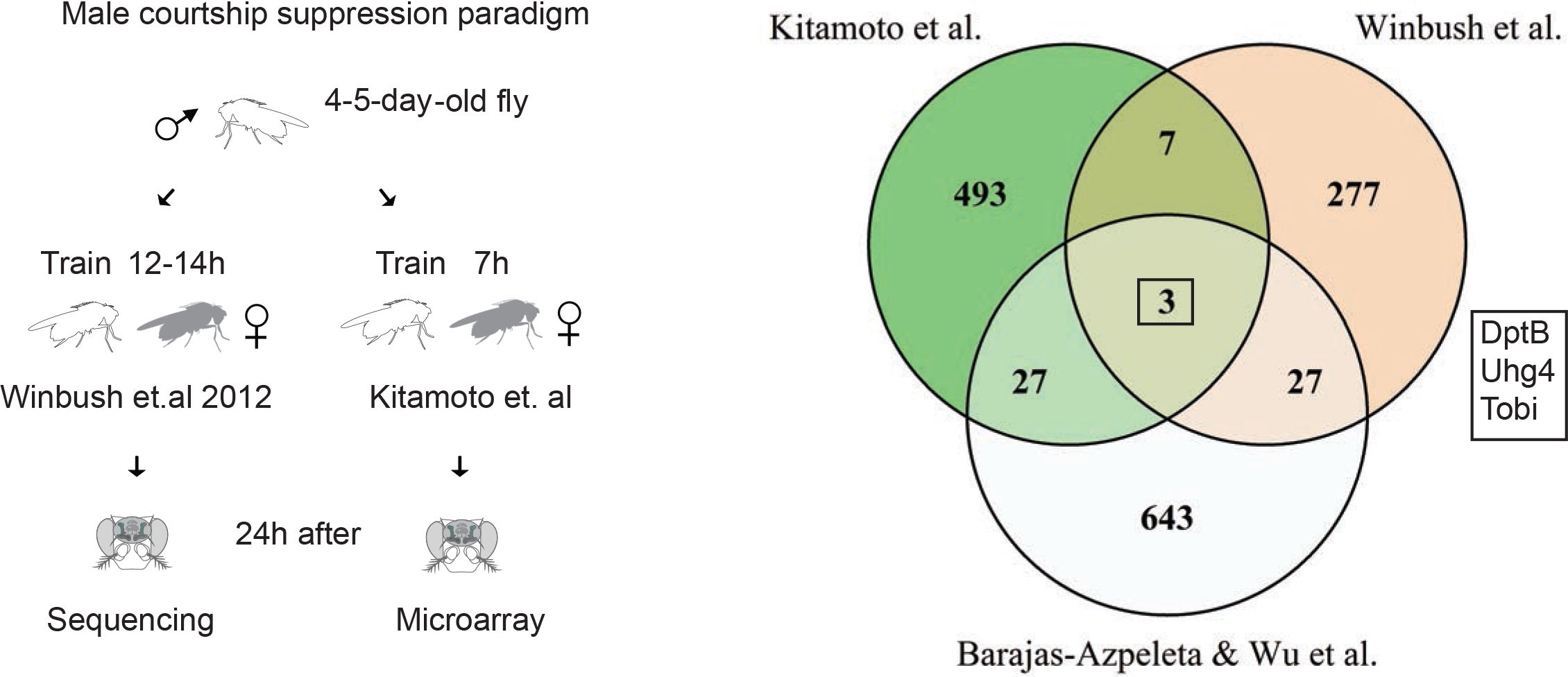
DptB expression changes after male courtship suppression training in various studies. Left, schematic representation of the male courtship suppression training used by other groups. Right, Venn diagram depicting gene expression changes in male courtship suppression paradigms performed by others, compared to this study. The common three genes are indicated in the box.

**Supplementary Figure S5.**
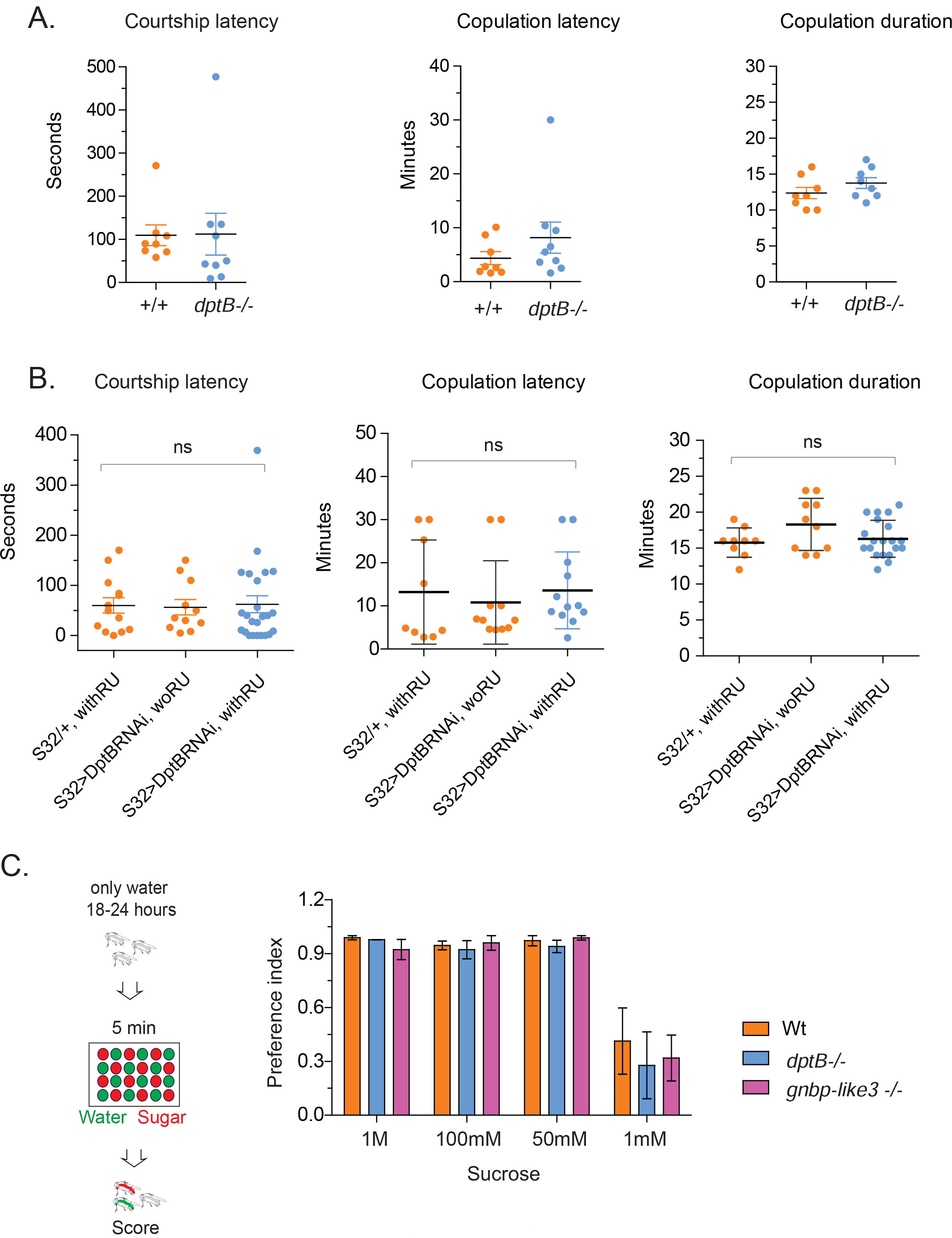
Innate behavior or operant learning is unchanged in DptB null or knock down flies. Left, courtship latency: after pairing time lag for the first display of courtship behavior; Middle, copulation latency: after pairing time lag before first successful mounting; Right, duration of copulation: once initiated, the duration of the first copulation; of DptB null flies (**A**) and DptB RNAi expressed in head fat body (**B**). (**C**) Detection of sucrose in different concentration is unaffected upon removal of DptB or GNBP-like3. Sucrose preference index of flies in indicated sucrose concentrations. The data are plotted as mean ± SEM. Ns stands for no statistically significant difference.

**Supplementary Figure S6.**
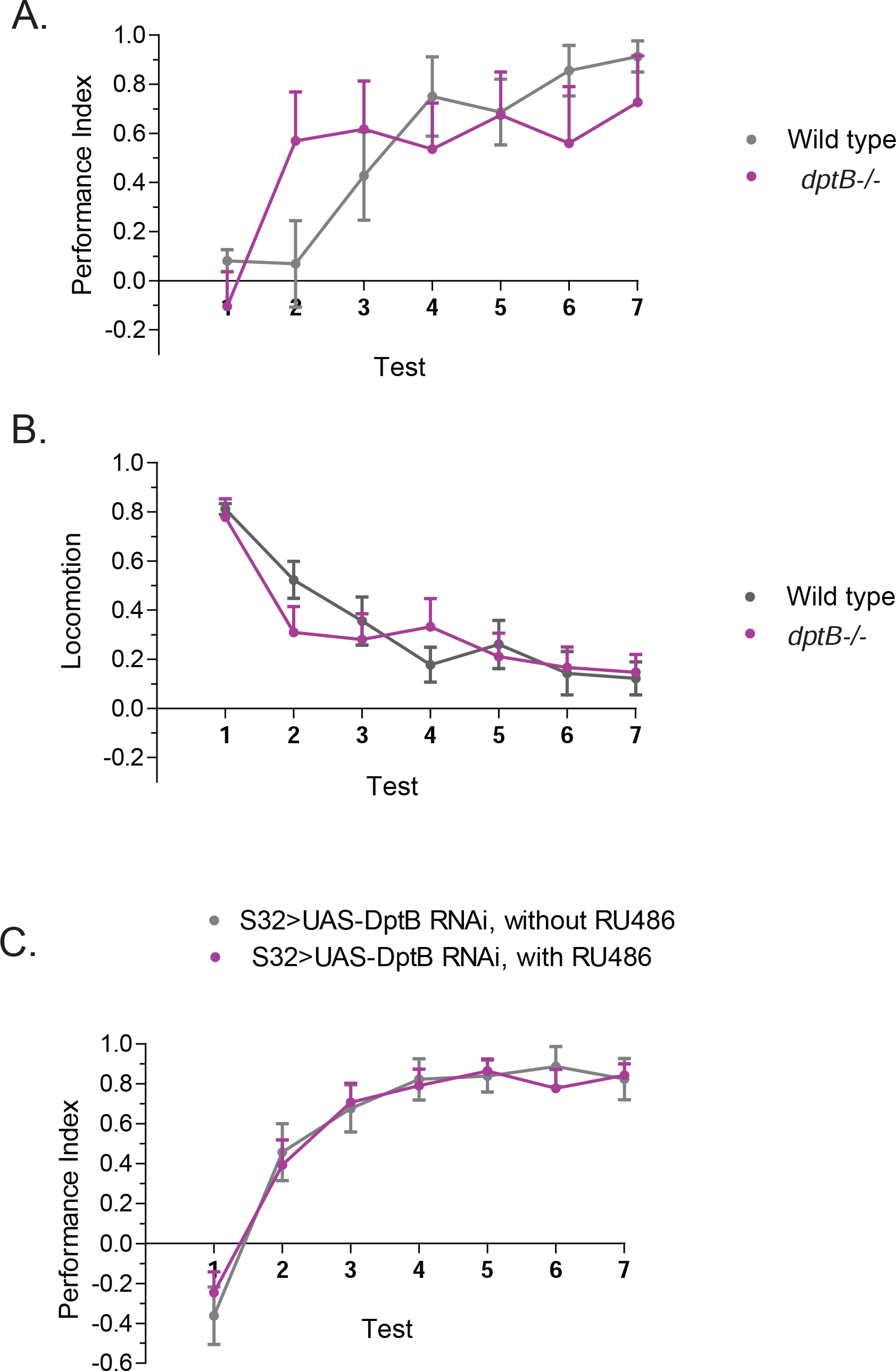
DptB knock down or knock out does not impair short-term aversive memory or locomotion. Memory is measured as the duration of the preferred response for unpunished side. Flies are either trained in a single trial protocol (1^st^) or multiple trial protocol where the single training protocol is repeated successively for 6 times [38,39]. (**A**) Removal of DptB gene or (**C**) expression of DptB RNAi in head fat body does not impair the ability to avoid the punished side. (**B**) The removal of DptB gene does not change locomotion. The decline in locomotion after each training is identical to that of wild type flies. The data are plotted as mean ± SEM.

**Supplementary Figure S7.**
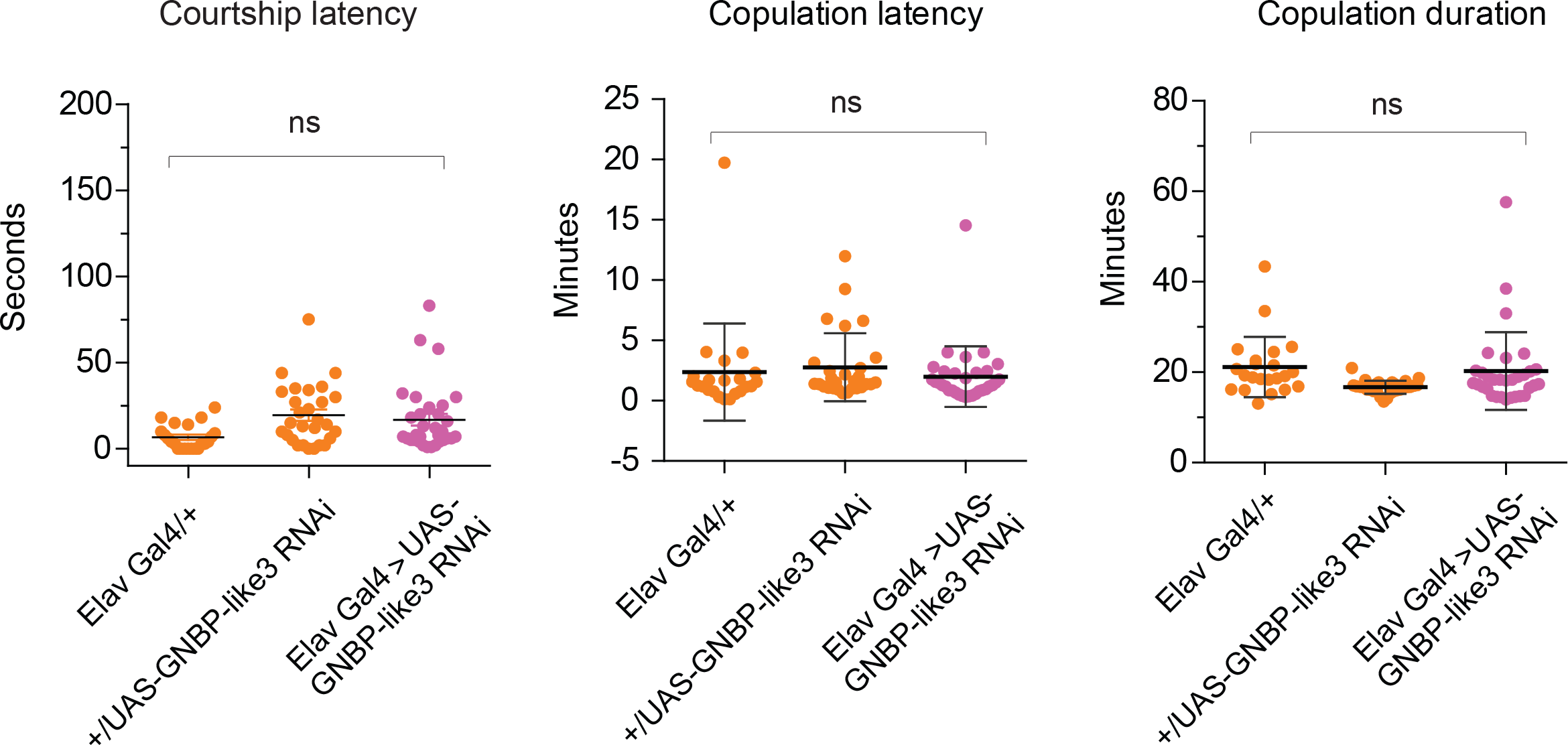
GNBP-like3 knock down does not impair innate courtship behavior. Left, courtship latency: after pairing time lag for the first display of courtship behavior; Middle, copulation latency: after pairing time lag before first successful mounting; Right, duration of copulation: once initiated, the duration of the first copulation.

**Supplementary Figure S8.**
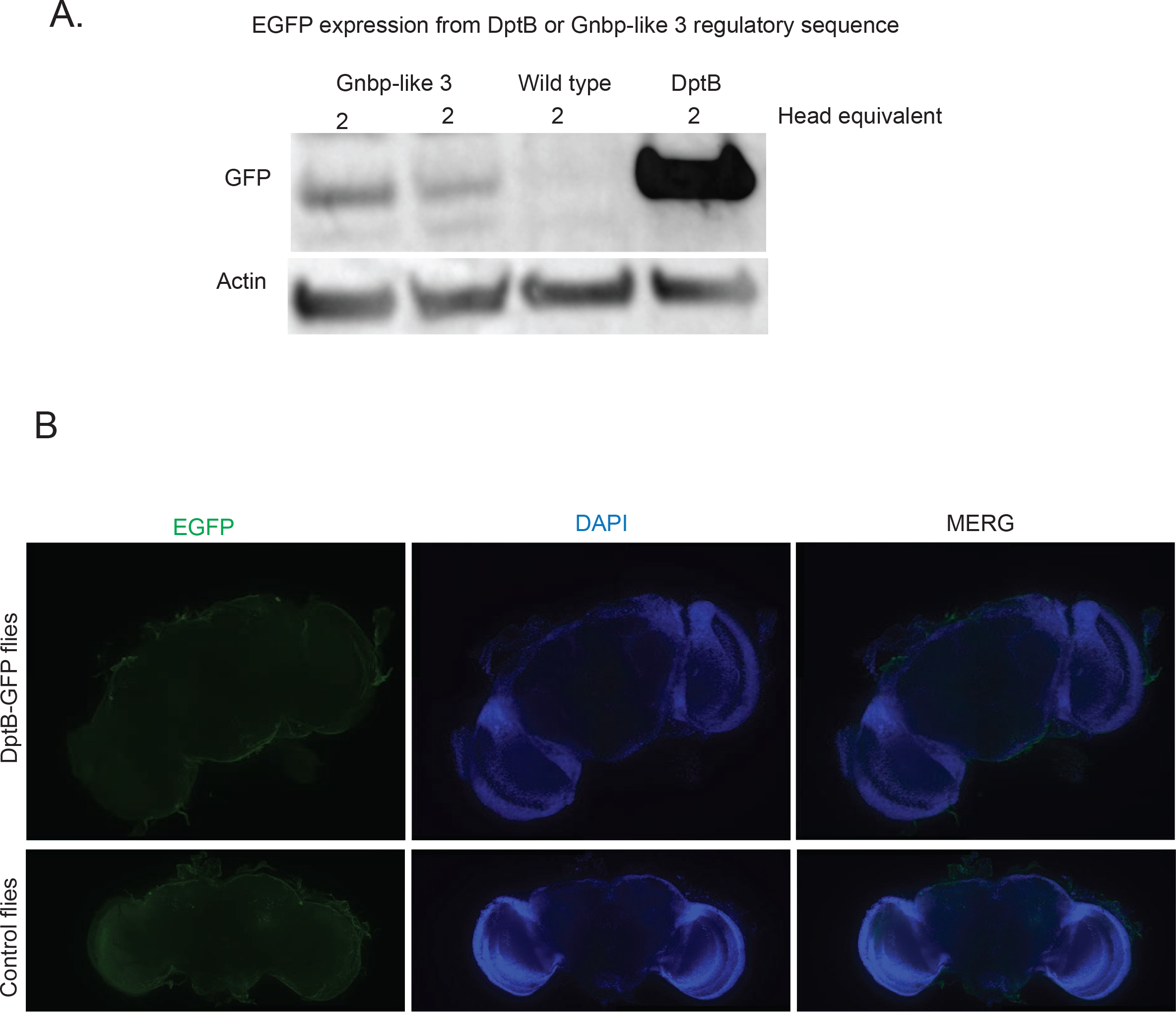
Expression of EGFP under GNBP-like 3 or DptB-regulatory sequences. (**A**) Western blot of fly heads expressing EGFP driven by GNBP-like3 or DptB regulatory sequences. The expression from GNBP-like3 locus is significantly lower than that from the DptB locus. The wild type fly head extracts serve as antibody specificity control. (B.) Expression of EGFP in central brain under DptB-regulatory sequence. Immunostaining of experimental (top), or control flies (bottom) for EGFP expression in the central brain region. There are no obvious differences between the EGFP and control flies, indicating DptB expression in the central brain is likely very low. Scale bar: 50 μm.

**Supplementary Figure S9.**
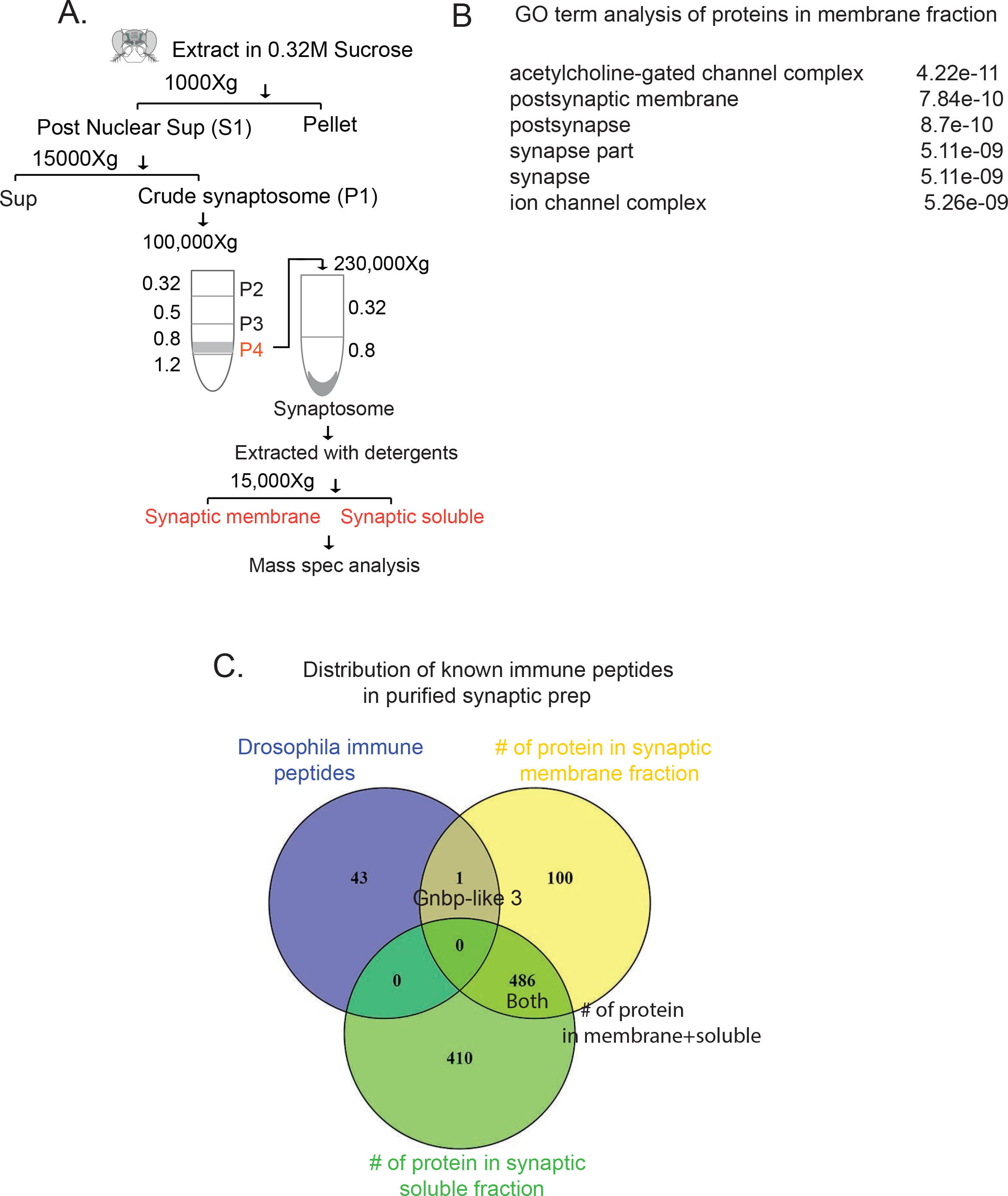
GNBP-like3 is present in the adult brain synaptosomes. (**A**). Schematic of the synaptosome purification. Heads were isolated from ~50 ml of flies and synaptosomes was prepared using a multistep procedure. (**B**) GO-term analysis of the 101 membrane proteins show top five GO-term categories are associated with synapse. (**C**) Venn diagram depicting distribution of immune peptides in the synaptosomes preparation. Among 43 annotated immune peptides only GNBP-like 3 is detected in the synaptosomes.

**Supplementary Figure S10.**
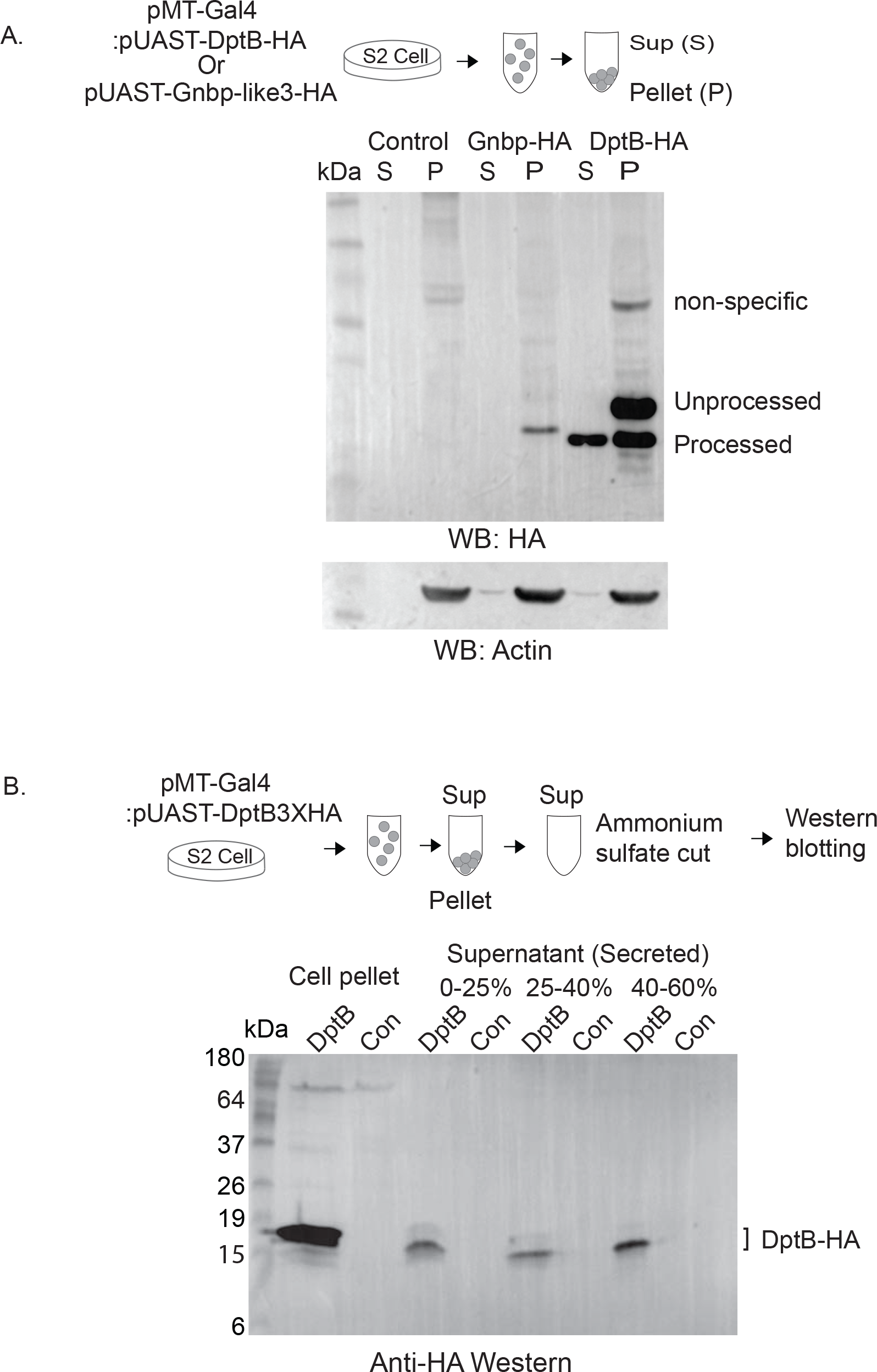
HA-tagged DptB is secreted in the media. (**A**) Top, schematic representation of the experiment. Bottom, 48h after transfection cells were harvested and using western blotting the media and the pellet were probed for secreted and total DptB or GNBP-like3 expression respectively. The media contains a smaller DptBHA fragment consistent with a processed and secreted peptide. The GNBP-like3 was detected only in the total cell lysates. Actin serves as a loading control. (**B**) Top, schematic representation of the experiment. Bottom, the media from DptB-HA-tagged stable cell line was harvested and subjected 0-25%, 25-40 % and 40-60% ammonium sulfate cut. Samples from each step were analyzed in western for HA-tagged DptB. Media from untransfected S2 cells were similarly treated and used as control (Con). Processed DptB is present in the media.

## References

1. Huh GS, Boulanger LM, Du H, Riquelme PA, Brotz TM, et al. (2000) Functional requirement for class I MHC in CNS development and plasticity. Science 290: 2155–2159.

2. Harris N, Braiser DJ, Dickman DK, Fetter RD, Tong A, et al. (2015) The Innate Immune Receptor PGRP-LC Controls Presynaptic Homeostatic Plasticity. Neuron 88: 1157–1164.

3. Stevens B, Allen NJ, Vazquez LE, Howell GR, Christopherson KS, et al. (2007) The classical complement cascade mediates CNS synapse elimination. Cell 131: 1164–1178.

4. Chen C, Itakura E, Nelson GM, Sheng M, Laurent P, et al. (2017) IL-17 is a neuromodulator of Caenorhabditis elegans sensory responses. Nature 542: 43–48.

5. Dissel S, Seugnet L, Thimgan MS, Silverman N, Angadi V, et al. (2015) Differential activation of immune factors in neurons and glia contribute to individual differences in resilience/vulnerability to sleep disruption. Brain Behav Immun 47: 75–85.

6. Bozler J, Kacsoh BZ, Chen H, Theurkauf WE, Weng Z, et al. (2017) A systems level approach to temporal expression dynamics in Drosophila reveals clusters of long term memory genes. PLoS Genet 13:e1007054.

7. McBride SM, Giuliani G, Choi C, Krause P, Correale D, et al. (1999) Mushroom body ablation impairs short-term memory and long-term memory of courtship conditioning in Drosophila melanogaster. Neuron 24: 967–977.

8. Majumdar A, Cesario WC, White-Grindley E, Jiang H, Ren F, et al. (2012) Critical role of amyloid-like oligomers of Drosophila Orb2 in the persistence of memory. Cell 148: 515–529.

9. Burke CJ, Waddell S (2011) Remembering nutrient quality of sugar in Drosophila. Curr Biol 21: 746–750.

10. McGinnis JP, Jiang H, Agha MA, Sanchez CP, Lange J, et al. (2016) Immediate perception of a reward is distinct from the reward’s long-term salience. Elife 5.

11. Dubnau J, Chiang AS, Grady L, Barditch J, Gossweiler S, et al. (2003) The staufen/pumilio pathway is involved in Drosophila long-term memory. Curr Biol 13: 286–296.

12. Lemaitre B, Hoffmann J (2007) The host defense of Drosophila melanogaster. Annu Rev Immunol 25: 697–743.

13. Lai Y, Gallo RL (2009) AMPed up immunity: how antimicrobial peptides have multiple roles in immune defense. Trends Immunol 30: 131–141.

14. Winbush A, Reed D, Chang PL, Nuzhdin SV, Lyons LC, et al. (2012) Identification of Gene Expression Changes Associated With Long-Term Memory of Courtship Rejection in Drosophila Males. G3: Genes|Genomes|Genetics 2: 1437–1445.

15. Wustmann G, Rein K, Wolf R, Heisenberg M (1996) A new paradigm for operant conditioning of Drosophila melanogaster. J Comp Physiol [A] 179: 429–436.

16. Lazareva AA, Roman G, Mattox W, Hardin PE, Dauwalder B (2007) A role for the adult fat body in Drosophila male courtship behavior. PLoS Genet 3: e16.

17. Fujii S, Amrein H (2002) Genes expressed in the Drosophila head reveal a role for fat cells in sex-specific physiology. EMBO J 21: 5353–5363.

18. Boulanger LM, Shatz CJ (2004) Immune signalling in neural development, synaptic plasticity and disease. Nat Rev Neurosci 5: 521–531.

19. Coulthard LG, Hawksworth OA, Woodruff TM (2018) Complement: The Emerging Architect of the Developing Brain. Trends Neurosci.

20. Widdowson EM, McCance RA (1935) The available carbohydrate of fruits: Determination of glucose, fructose, sucrose and starch. Biochem J 29: 151–156.

21. Sánchez-Castillo CP, Dewey PJ, Lara JJ, Henderson DL, de Lourdes Solano Ma, et al. (2000) The starch and sugar content of some Mexican cereals, cereal products, pulses, snack foods, fruits and vegetables. Journal of Food Composition and Analysis 13: 157–170.

22. Huang Z, Kingsolver MB, Avadhanula V, Hardy RW (2013) An antiviral role for antimicrobial peptides during the arthropod response to alphavirus replication. J Virol 87: 4272–4280.

23. De Gregorio E, Spellman PT, Rubin GM, Lemaitre B (2001) Genome-wide analysis of the Drosophila immune response by using oligonucleotide microarrays. Proc Natl Acad Sci U S A 98: 12590–12595.

24. Irving P, Troxler L, Heuer TS, Belvin M, Kopczynski C, et al. (2001) A genome-wide analysis of immune responses in Drosophila. Proc Natl Acad Sci U S A 98: 15119–15124.

25. Candille SI, Kaelin CB, Cattanach BM, Yu B, Thompson DA, et al. (2007) A-defensin mutation causes black coat color in domestic dogs. Science 318: 1418–1423.

26. Brogden KA, Guthmiller JM, Salzet M, Zasloff M (2005) The nervous system and innate immunity: the neuropeptide connection. Nat Immunol 6: 558–564.

27. Cao Y, Chtarbanova S, Petersen AJ, Ganetzky B (2013) Dnr1 mutations cause neurodegeneration in Drosophila by activating the innate immune response in the brain. Proc Natl Acad Sci U S A 110: E1752–1760.

28. Soscia SJ, Kirby JE, Washicosky KJ, Tucker SM, Ingelsson M, et al. (2010) The Alzheimer’s disease-associated amyloid beta-protein is an antimicrobial peptide. PLoS One 5: e9505.

29. Williams WM, Castellani RJ, Weinberg A, Perry G, Smith MA (2012) Do beta-defensins and other antimicrobial peptides play a role in neuroimmune function and neurodegeneration? Scientific World Journal 2012: 905785.

30. Trapnell C, Williams BA, Pertea G, Mortazavi A, Kwan G, et al. (2010) Transcript assembly and quantification by RNA-Seq reveals unannotated transcripts and isoform switching during cell differentiation. Nat Biotechnol 28: 511–515.

31. Gill J, Park Y, McGinnis JP, Perez-Sanchez C, Blanchette M, et al. (2017) Regulated Intron Removal Integrates Motivational State and Experience. Cell 169: 836–848 e815.

32. Li L, Sanchez CP, Slaughter BD, Zhao Y, Khan MR, et al. (2016) A Putative Biochemical Engram of Long-Term Memory. Curr Biol 26: 3143–3156.

33. Helaine S, Thompson JA, Watson KG, Liu M, Boyle C, et al. (2010) Dynamics of intracellular bacterial replication at the single cell level. Proc Natl Acad Sci U S A 107: 3746–3751.

34. Sakai T, Kitamoto T (2006) Differential roles of two major brain structures, mushroom bodies and central complex, for Drosophila male courtship behavior. J Neurobiol 66: 821–834.

35. Winbush A, Reed D, Chang PL, Nuzhdin SV, Lyons LC, et al. (2012) Identification of gene expression changes associated with long-term memory of courtship rejection in Drosophila males. G3 (Bethesda) 2: 1437–1445.

36. Benjamini Y, and Hochberg Y. (1995) Controlling the false discovery rate: a practical and powerful approach to multiple testing. Journal of the Royal Statistical Society Series 57: 289–300.

37. Robinson MD, McCarthy DJ, Smyth GK (2010) edgeR: a Bioconductor package for differential expression analysis of digital gene expression data. Bioinformatics 26: 139–140.

38. Diegelmann S, Zars M, Zars T (2006) Genetic dissociation of acquisition and memory strength in the heat-box spatial learning paradigm in Drosophila. Learn Mem 13: 72–83.

39. Putz G, Heisenberg M (2002) Memories in drosophila heat-box learning. Learn Mem 9: 349–359.

